# Control of Ca^2+^ signals by astrocyte nanoscale morphology at tripartite synapses

**DOI:** 10.1101/2021.02.24.432635

**Authors:** Audrey Denizot, Misa Arizono, Valentin U. Nägerl, Hugues Berry, Erik De Schutter

## Abstract

Much of the Ca^2+^ activity in astrocytes is spatially restricted to microdomains and occurs in fine processes that form a complex anatomical meshwork, the so-called spongiform domain. A growing body of literature indicates that those astrocytic Ca^2+^ signals can influence the activity of neuronal synapses and thus tune the flow of information through neuronal circuits. Because of technical difficulties in accessing the small spatial scale involved, the role of astrocyte morphology on Ca^2+^ microdomain activity remains poorly understood. Here, we use computational tools and idealized 3D geometries of fine processes based on recent super-resolution microscopy data to investigate the mechanistic link between astrocytic nanoscale morphology and local Ca^2+^ activity. Simulations demonstrate that the nano-morphology of astrocytic processes powerfully shapes the spatio-temporal properties of Ca^2+^ signals and promotes local Ca^2+^ activity. The model predicts that this effect is attenuated upon astrocytic swelling, hallmark of brain diseases, which we confirm experimentally in hypo-osmotic conditions. Upon repeated neurotransmitter release events, the model predicts that swelling hinders astrocytic signal propagation. Overall, this study highlights the influence of the complex morphology of astrocytes at the nanoscale and its remodeling in pathological conditions on neuron-astrocyte communication at so-called tripartite synapses, where astrocytic processes come into close contact with pre- and postsynaptic structures.

**Table of Contents:** *Main Points:*

- Astrocyte nano-morphology favors the compartmentalization of biochemical signals
- This compartmentalization promotes local Ca^2+^ activity and signal propagation robustness
- In contrast, its pathological remodeling upon swelling attenuates Ca^2+^ activity

*Table of Contents Image:* Figure 1:
Proposed mechanisms that regulate astrocytic Ca^2+^ activity in perisynaptic astrocytic processes.
Our simulation results demonstrate that the nano-morphology of astrocytic processes, consisting in the alternation of nodes and shafts, favors the compartmentalization of biochemical signals. This compartmentalization promotes local Ca^2+^ activity and signal propagation robustness. Astrocyte swelling, observed in pathological conditions such as brain injury, stroke and epilepsy, results in an increased shaft width without altering node size. Our results suggest that such pathological alterations of the nanoscale morphology of astrocytes result in a decreased local Ca^2+^ activity, which we confirm experimentally in hypo-osmotic conditions. Upon repeated neuronal stimuli, we predict that swelling hinders astrocytic signal propagation. Overall, this study highlights the impact of astrocyte nano-morphology on astrocyte activity at tripartite synapses, in health and disease.

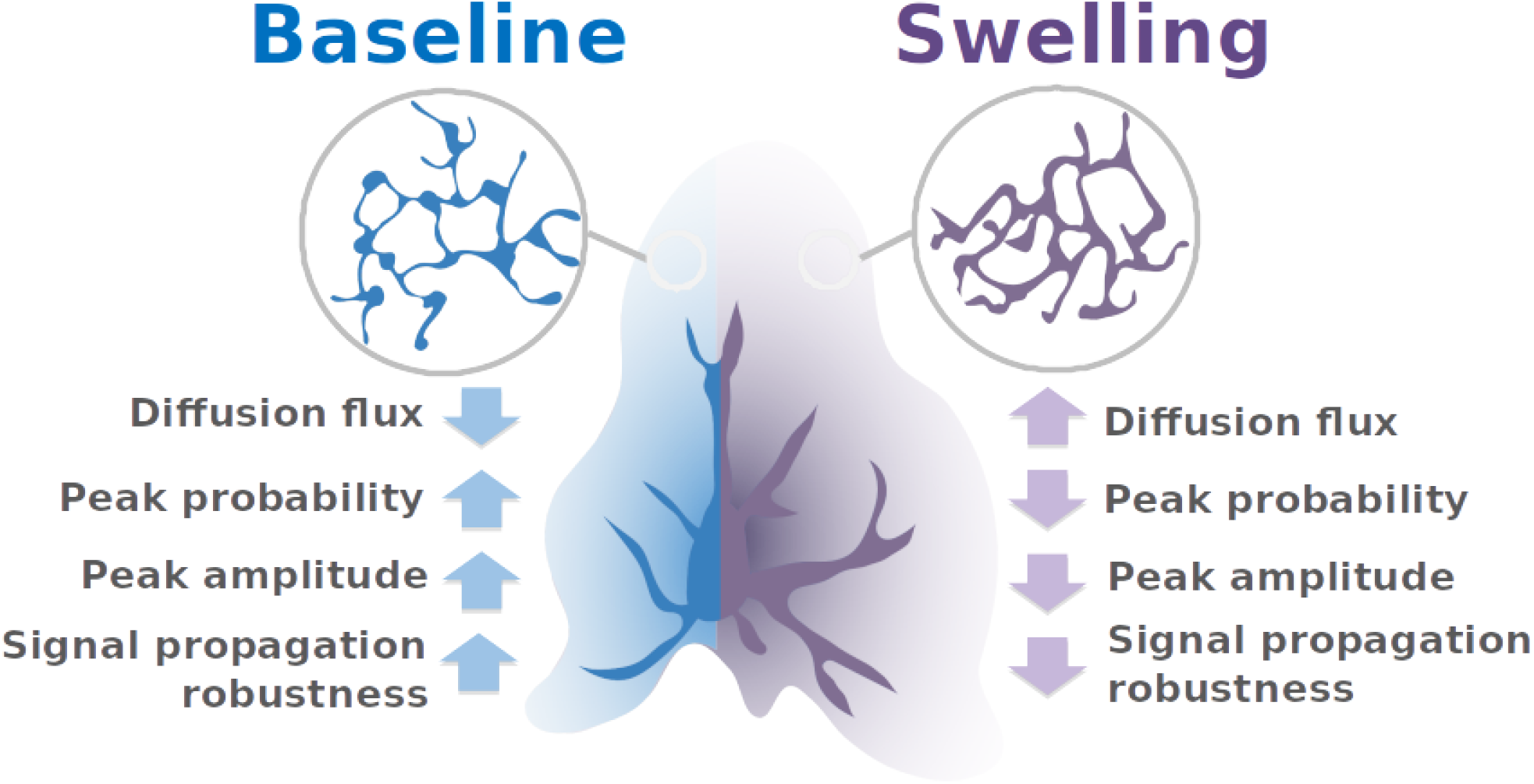

## Introduction

Astrocytes are glial cells of the central nervous system that are essential for brain development and function Verkhratsky & Nedergaard (2018). They notably modulate neuronal communication at synapses. Astrocytic Ca^2+^ signals are triggered by neurotransmitters released by active neurons, which can trigger the release of neuroactive molecules by the astrocyte, referred to as gliotransmitters. The first type of astrocytic Ca^2+^ signals that has been observed was Ca^2+^ waves that propagate through gap junctions in astrocyte networks Giaume & Venance (1998). Ca^2+^ waves have also been observed in the branches of single astrocytes, sporadically propagating to the soma Haustein et al. (2014); Bindocci et al. (2017). The recent development of Ca^2+^ imaging techniques with improved spatial and temporal resolution has revealed the existence of spatially-restricted Ca^2+^ signals in astrocytes, referred to as microdomains or hotspots Di Castro et al. (2011); Panatier et al. (2011); Stobart, Ferrari, Barrett, Glück, et al. (2018); Srinivasan et al. (2015); Shigetomi et al. (2013); Sherwood et al. (2017); Otsu et al. (2015); Lind et al. (2013); Bindocci et al. (2017); Agarwal et al. (2017); Arizono et al. (2020); Otsu et al. (2015). These local Ca^2+^ signals account for the vast majority (≈ 80 %) of astrocytic Ca^2+^ activity and occur in fine processes, which occupy 75 % of the astrocytic volume Bindocci et al. (2017), forming the spongiform domain, also referred to as the gliapil. Given that one astrocyte may contact tens of thousands of synapses simultaneously Bushong et al. (2002) via these fine processes, local and fast Ca^2+^ signals might enable the astrocyte to powerfully yet precisely control the flow of information through synaptic circuits. Importantly, reactive astrocytes, hallmark of brain diseases, display aberrant amplitude, duration, frequency and spatial spread of Ca^2+^ signals Shigetomi et al. (2019); Nedergaard et al. (2010); Lee et al. (2022).

Cellular micro-morphology lends itself to the compartmentalization of biochemical signals. For example, the anatomical design of dendritic spines restricts the diffusion of Ca^2+^ to the activated synapse, which reduces cross-talk between nearby synapses Santamaria et al. (2011); Tonnesen et al. (2014); Yuste et al. (2000); Noguchi et al. (2005); Yasuda (2017); Holcman & Schuss (2011). The complex shapes of Bergmann glia Grosche et al. (1999) and perisynaptic “astrocytic compartments” along major branches Panatier et al. (2011) have been proposed to restrict Ca^2+^ signals to the vicinity of synapses. Fine processes of the spongiform domain, however, cannot be resolved by diffraction-limited light microscopy Rusakov (2015), so that the contribution of their morphology to shaping local Ca^2+^ signals is poorly understood. Our recent 3D STED study Arizono et al. (2020) revealed the structural basis of compartmentalized spontaneous Ca^2+^ signals in fine astrocytic processes. Importantly, pathological changes in astrocytic morphology Lafrenaye & Simard (2019), such as “astrocytic swelling”, are paired with aberrant Ca^2+^ signals Shigetomi et al. (2019). We have recently reported that swelling can also occur at the level of fine astrocytic processes Arizono, Bancelin, et al. (2021). The effect of such a remodeling of astrocytic nano-morphology on the local Ca^2+^ signals involved in regulating synapses remains yet unclear. Together, there is a great interest in understanding the mechanistic link between the nano-morphology of astrocytic processes and Ca^2+^ profiles in health and disease. Computational approaches make it possible to simulate different geometrical scenarios, in a much more systematic and controlled way than what could be done experimentally. Computational models can thus help us gain insights into the impact of morphological parameters on Ca^2+^ activity.

Here, we use computational tools to explore the role of the anatomical design of the gliapil. To do so, we perform simulations in branchlet geometries that we designed based on super-resolution microscopy data reported in live tissue Arizono et al. (2020), consisting of nodes, which host Ca^2+^ microdomains, and their intervening shafts of variable widths. Our simulation results suggest that the nanoscale design of the spongiform domain effectively decreases diffusion flux, which increases Ca^2+^ peak probability, duration and amplitude in the stimulated and neighboring nodes. To test those predictions, we performed Ca^2+^ recordings in organotypic hipppocampal cultures in hypo-osmotic conditions, where the normal node-shaft arrangement is altered. In line with our model predictions, Ca^2+^ activity in hypo-osmotic conditions was decreased compared to Ca^2+^ activity in normal tissue. We further found that, upon repeated neuronal stimulation, thin shafts allow signal propagation even if some stimuli are omitted, thus allowing for a more robust signal propagation.

Overall, our study sheds light on the influence of the nanoscale morphology of the complex spongiform domain of astrocytes on Ca^2+^ microdomain activity and indicates that pathological morphological changes may substantially affect their Ca^2+^ activity.

## Methods

### Stochastic spatially-explicit voxel-based simulations

In order to model astrocyte Ca^2+^ signals in astrocyte branchlets, we have used the voxel-based “GCaMP” implementation of the Inositol 3-Phosphate (IP_3_) receptor-dependent Ca^2+^ signaling model from Denizot et al Denizot et al. (2019), using the same reaction scheme and parameter values (Fig 2B). Briefly, we model Ca^2+^ fluxes in and out of the cytosol, mediated by Ca^2+^ channels and pumps on the endoplasmic reticulum (ER) and on the plasma membrane. Ca^2+^ signals occur when some IP_3_R channels are in the open state. IP_3_ can be synthesized by the Ca^2+^-dependent activity of phospholipase C *δ* (PLC*δ*) and the removal of IP_3_ molecules from the cytosol is expressed as a single decay rate. IP_3_R kinetics is described by a Markov model, derived from De Young & Keizer’s model De Young & Keizer (1992). Each IP_3_R molecule contains one IP_3_ binding site and two Ca^2+^ binding sites. An IP_3_R is in the open state when in state {110} (first Ca site and IP_3_ bound, second Ca site free). Depending on the simulation, other diffusing molecules were added to the model, such as the fluorescent molecule ZsGreen and fluorescent Ca^2+^ indicators, here 10 *µ*M of GCaMP6s. GCaMPs are genetically-encoded Ca^2+^ indicators (GECIs) that are derived from the fluorescent protein GFP and the Ca^2+^ buffer calmodulin (see Shigetomi et al. (2016) for a review on GECIs). For further details on the kinetic scheme and model assumptions, please refer to Denizot et al. 2019 Denizot et al. (2019).

**Figure 2:**
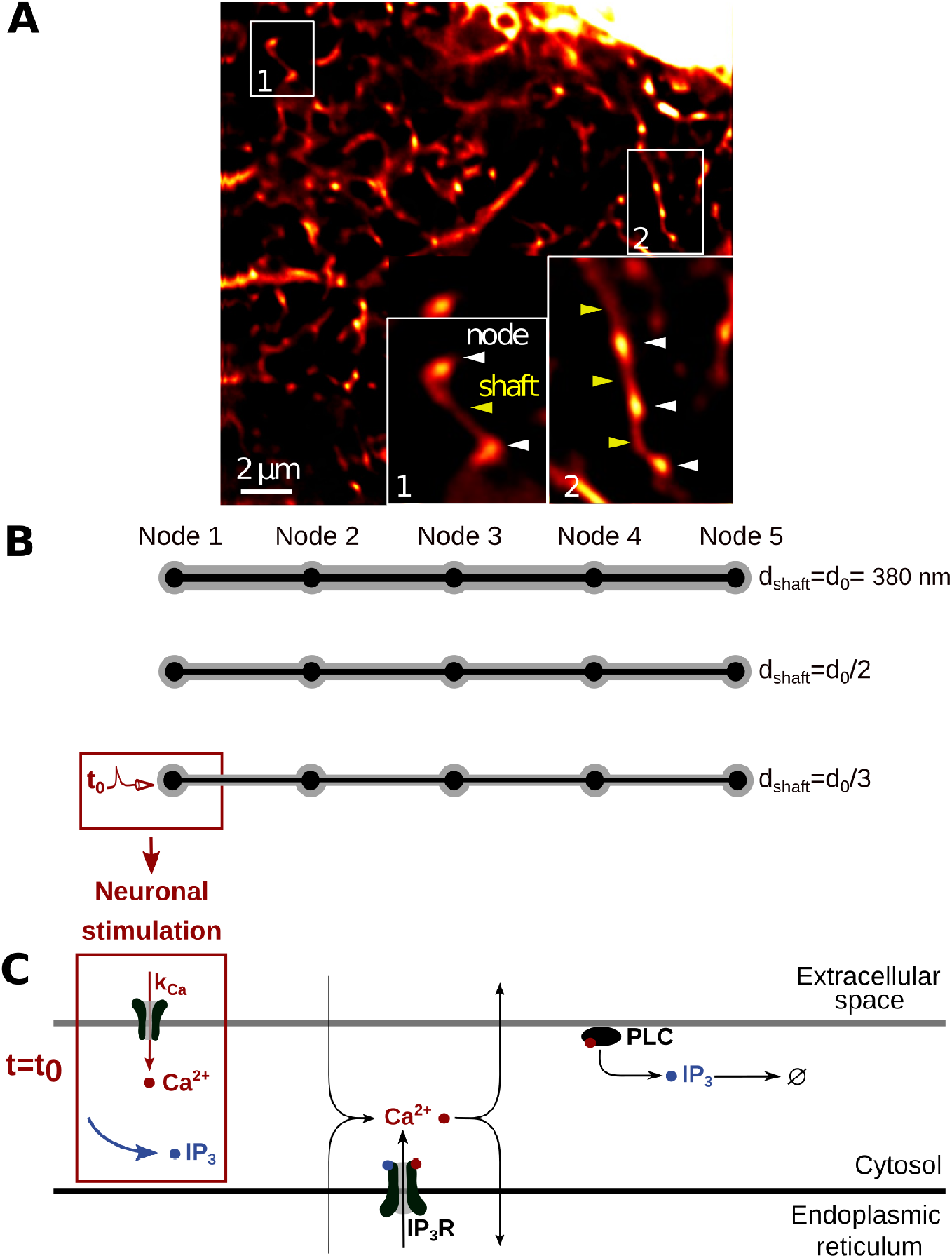
Geometries and kinetic scheme used for simulating Ca^2+^ dynamics in node/shaft structures of the gliapil. (*A*) Representative STED image showing the astrocytic spongiform domain. Zoom-in images show its anatomical units: nodes and shafts. (*B*) Geometries reproducing node/shaft geometries of the gliapil Arizono et al. (2020); Panatier et al. (2014) were designed. Nodes are approximated as spheres of diameter 380 nm and shafts as 1 *µ*m-long cylinders. The geometries designed in this study, referred to as “5 nodes”, contain 5 identical nodes and 4 identical shafts. Unless specified otherwise, ER geometry (black) also consists in node/shaft successions (see Methods). Geometries were characterized by different shaft widths: *d*_shaft_ = *d*_0_ = 380 nm, 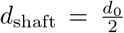 and 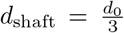. The associated cytosolic volume, plasma and ER membrane areas are presented in Table 1. (*C*) Biochemical processes included in the model. Ca^2+^ can enter/exit the cytosol from/to the extracellular space or the endoplasmic reticulum (ER), resulting from the activity of Ca^2+^ channels/pumps. Ca^2+^ and IP_3_ diffuse in the cytosol following Brownian motion. The kinetics of *IP*_3_*R* channels corresponds to the 8-state Markov model from Denizot et al. (2019), adapted from De Young & Keizer (1992); Bezprozvanny et al. (1991). When both IP_3_ and Ca^2+^ are bound to IP_3_R activating binding sites, the IP_3_R is in open state and Ca^2+^ enters the cytosol. Ca^2+^ can activate Phospholipase C *δ* (PLC*δ*), which results in the production of IP_3_. For more details, please refer to Denizot et al. (2019). Neuronal stimulation is simulated as an infusion of IP_3_ in the cytosol and the opening of Ca^2+^ channels at the plasma membrane with an influx rate *k*_*Ca*_ (see Methods).

The model was implemented using STEPS (http://steps.sourceforge.net/), a python package performing exact stochastic simulation of reaction-diffusion systems Hepburn et al. (2012). More presicely, STEPS uses a spatialized implementation of Gillespie’s SSA algorithm Gillespie (1977); Isaacson & Isaacson (2009); Smith & Grima (2018). Simulations in STEPS can be performed in complex geometries in 3 spatial dimensions. Space is divided into well-mixed tetrahedral compartments, referred to as voxels. Reactions between 2 molecules can only occur if they are located within the same voxel. Diffusion events are modeled as a decrease of the number of molecules in the original voxel and an increase in the number of molecules in its neighboring voxel. Boundary conditions, except when specified otherwise, were reflective. In other words, mobile molecules could not diffuse away from the geometry, as if they were “bouncing” onto the plasma membrane. STEPS enables to compute, in complex 3D geometries, reactions and diffusion in the cytosol as well as reactions between cytosolic molecules and molecules located at the plasma or ER membrane.

### Geometries

Typical astrocyte branchlet geometries were designed from their recent experimental characterization in live tissue at high spatial resolution (50 nm in x-y) Arizono et al. (2020). Those geometries consist in alternations of bulbous structures, nodes, connected to each other with cylindrical structures, shafts. Geometries with different shaft widths d_shaft_ were designed using Trelis software (https://www.csimsoft.com/trelis, Fig 2A). The geometry of a node was approximated as being a sphere of diameter 380 nm. Shaft geometry consisted in a 1*µ*m long cylinder. Shaft diameter was defined relative to node diameter. For example, shaft diameter was the same as node diameter, i.e d_shaft_ =*d*_0_=380 nm. Similarly, shaft diameter was 190 nm and 127 nm for 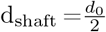 and 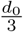, respectively. Cones were positioned between spheres and cylinders in order to create a smoother transition between nodes and shafts, better approximating the geometry observed experimentally. Cytosolic volume was thus *V*_1_=0.620 *µm*^3^, *V*_2_=0.263 *µm*^3^ and *V*_3_=0.195 *µm*^3^, for 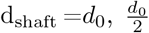 and 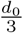, respectively. A subset of simulations was performed in a geometry with *V*_1_=0.258 *µm*^3^. This geometry is characterized, similarly to geometries with d_shaft_ =*d*_0_, by a node/shaft width ratio of 1. It contains cylinders of length 750 nm, diameter 285 nm and spheres of diameter 285 nm. As a first approximation, ER geometry was considered to be similar to the geometry of the astrocyte branchlet: node/shaft successions. ER nodes were aligned with cytosolic nodes. As no quantification of the ratio between astrocytic ER volume and cellular volume was found in the literature, ER volume was 10% of the total branchlet volume, based on available data in neurons Spacek & Harris (1997). As the shape and distribution of the ER in fine processes have not been characterized in live tissue but are likely highly variable, additional simulations were performed in meshes with various ER shapes: “No ER”, “Node ER” and “Cyl ER”, in which there was no ER, discontinuous ER in nodes or cylindrical ER, respectively (Fig S7-S9). The cytosolic volume, plasma and ER membrane surface areas of those 3D geometries are presented in Table 1.

**Table 1:** Characteristics of the geometries of astrocyte branchlets used in this study. *V*_cyt_ is the cytosolic volume, *S*_PM_ is the area of the plasma membrane and *S*_ER_ is the area of the ER membrane. Volumes are expressed in *nm*^3^ and areas in *nm*^2^. Meshes are available at http://modeldb.yale.edu/266928, access code: lto42@tpk3D?

| Geom | $V_{\text{cyt}} (\text{nm}^3)$ | $S_{\text{PM}} (\text{nm}^2)$ | $S_{\text{ER}} (\text{nm}^2)$ |
| --- | --- | --- | --- |
| “5nodes” $d_{\text{shaft}} = d_0$ | $6.20 \times 10^8$ | $7.73 \times 10^6$ | $2.19 \times 10^6$ |
| “5nodes” $d_{\text{shaft}} = d_0/2$ | $2.63 \times 10^8$ | $4.95 \times 10^6$ | $1.28 \times 10^6$ |
| “5nodes” $d_{\text{shaft}} = d_0/3$ | $1.95 \times 10^8$ | $4.10 \times 10^6$ | $9.99 \times 10^5$ |
| “No ER” $d_{\text{shaft}} = d_0$ | $6.73 \times 10^8$ | $7.73 \times 10^6$ | 0.00 |
| “No ER” $d_{\text{shaft}} = d_0/2$ | $2.83 \times 10^8$ | $4.95 \times 10^6$ | 0.00 |
| “No ER” $d_{\text{shaft}} = d_0/3$ | $2.10 \times 10^8$ | $4.09 \times 10^6$ | 0.00 |
| “Node ER” $d_{\text{shaft}} = d_0$ | $6.67 \times 10^8$ | $7.75 \times 10^6$ | $4.17 \times 10^5$ |
| “Node ER” $d_{\text{shaft}} = d_0/2$ | $2.74 \times 10^8$ | $4.96 \times 10^6$ | $4.37 \times 10^5$ |
| “Node ER” $d_{\text{shaft}} = d_0/3$ | $2.00 \times 10^8$ | $4.11 \times 10^6$ | $4.41 \times 10^5$ |
| “Cyl ER” $d_{\text{shaft}} = d_0$ | $6.27 \times 10^8$ | $7.74 \times 10^6$ | $2.03 \times 10^6$ |
| “Cyl ER” $d_{\text{shaft}} = d_0/2$ | $2.77 \times 10^8$ | $4.95 \times 10^6$ | $8.78 \times 10^5$ |
| “Cyl ER” $d_{\text{shaft}} = d_0/3$ | $2.07 \times 10^8$ | $4.09 \times 10^6$ | $5.86 \times 10^5$ |

A sensitivity study was performed to investigate the effect of voxel size on the kinetics of the molecular interactions modeled. Information on the voxel sizes of the different meshes used is presented in Table S1. Results are presented in Figure S1. Meshes that contained voxels that were *<* 50 nm^3^ were characterized by aberrant kinetics, resulting in aberrant average numbers of molecules in a given state. Those results thus suggest that to prevent errors due to voxel size, meshes should not display voxel sizes that are*<* 50 nm^3^. We have thus made sure, while meshing the geometries in which simulations were ran, that no voxels were *<* 50 nm^3^. Minimum voxel size was 443 nm^3^, 1100 nm^3^ and 447 nm^3^, for 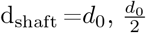 and 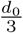 geometries, respectively.

In a subset of simulations, ER geometry varied. The shape of the cell was the same as in “5nodes” geometries (Fig 2). ER geometry consisting of node/shaft alternations, described above, is referred to as “Node/shaft ER”. “No ER” geometry contains no ER. “Node ER” is characterized by a discontinuous ER geometry, consisting in spheres of diameter 54 nm, located in cellular nodes. “Cyl ER” corresponds to a cylindrical ER, of length *l*_ER_= 6274 nm and a diameter of 108, 54 and 36 nm, for 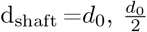 and 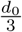, respectively. The associated cytosolic volume, ER and plasma membrane area are presented in Table 1.

### Protocol for simulating bleaching experiments

In order to test whether the idealized geometries presented in Fig 2 are a good approximation of the spongiform ultrastructure of astrocyte branchlets, we have simulated their fluorescence recovery after photobleaching (FRAP) experiments. Briefly, laser pulses are simulated on a node (region of interest) while the fluorescence level is being recorded. At bleaching time, the fluorescence level in the region of interest decreases to *I*_0_. Then, because of the diffusion of fluorescent molecules into the region of interest, fluorescence increases until it reaches a new steady state, *I*_inf_. We characterize node compartmentalization by measuring the time *τ* taken by fluorescing molecules to diffuse into the node to reach *I*_inf_. In other words, a high node compartmentalization will be associated with a high value of *τ*. Thus, 3 main parameters characterize bleaching traces: *I*_0_, *τ* and *I*_inf_.

To mimic bleaching experiments in fine branchlets performed by Arizono et al Arizono et al. (2020), Zs-Green molecules were added to simulation space. After 2 seconds of simulation, providing the basal level of fluorescence, 60% of ZsGreen molecules were bleached. In order to fit *I*_0_ and *I*_inf_ that were measured experimentally, and as bleaching time lasted 10 ms in experiments and 1 ms in simulations, the bleached volume in simulations was adjusted depending on the geometry (see Fig 3A). Bleaching was simulated as a transition from ZsGreen molecules to ZsGreen-bleached molecules, the latter being considered as non-fluorescing molecules. Screenshots of simulations, illustrating the diffusion of ZsGreen and ZsGreen-bleached molecules, are presented in Fig S2B. The number of ZsGreen molecules in the central node was recorded over simulation time and a fit was performed following equation 1 to determine the values of *I*_0_, *I*_inf_ and *τ*.

**Figure 3:**
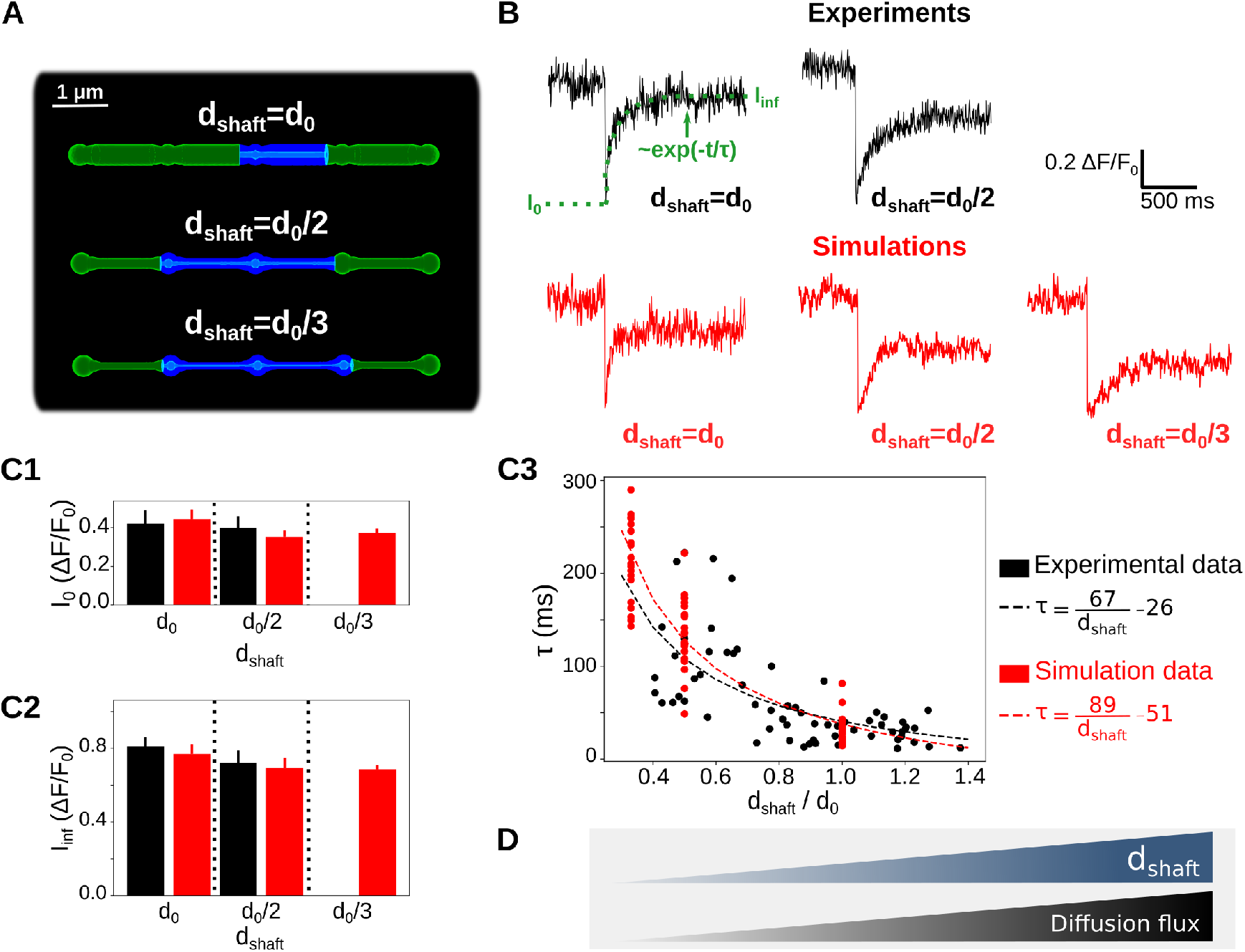
Simulations confirm that thin shafts favor node compartmentalization. (*A*) Geometries of different shaft widths 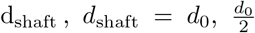 and 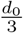, used in the bleaching simulations. Blue color represents the bleached volume, which varied depending on the value of d_shaft_ in order to fit experimental values of *I*_0_ and *I*_*inf*_. (*B*) Representative experimental (top) and simulation (bottom) traces for different shaft width values. *I*_0_, *I*_*inf*_ and *τ* were calculated using Eq 1. Note that simulations were also performed for 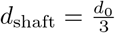, (*C*) Quantification of *I*_0_ (C1), *I*_*inf*_ (C2) and *τ* (C3) values in simulations (red) compared to experiments (black). Note that no experimental data was available for 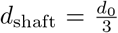. In C1 and C2, n=5×2 and 20×3 for experiments and simulations, respectively. Data are presented as mean ± STD. In C3, n=66 and n=20×3 for experiments and simulations, respectively. *τ* is negatively correlated to d_shaft_ in experiments (n=66 from 7 slices; Spearman r=-0.72, p*<*0.001 ***) and simulations (n=60; Spearman r=-0.89, p*<*0.001 ***). Black and red lines represent curve fit of *τ* as a function of d_shaft_ of the form 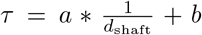 for experiments and simulations, respectively. (*D*) Schematic summarizing the conclusion of this figure: diffusion flux increases with d_shaft_. In that sense, thin shafts favor node compartmentalization. Data in panels C1 and C2 are represented as mean ± STD, n=20 for each geometry.

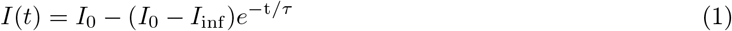

where I(t) is the level of fluorescence measured at time t. The coefficient of diffusion, *D*_ZsGreen_, and the concentration, [ZsGreen], of ZsGreen were adjusted to fit experimental data. Indeed, the amplitude of [ZsGreen] fluctuations at steady state is inversely proportional to the number of ZsGreen molecules in the geometry. In other words, fluorescence signals are more noisy when [ZsGreen] is low. Moreover, the autocorrelation of those fluctuations depends on the coefficient of diffusion of ZsGreen, *D*_ZsGreen_. If *D*_ZsGreen_ increases, the autocorrelation of Lag, where Lag is the autocorrelation delay, will decrease faster as Lag increases. Comparing the fluctuations of [ZsGreen] and its autocorrelation in experiments and in simulations thus enabled to find the values of *D*_ZsGreen_ and of [ZsGreen] that allowed for the best fit to experimental data. In the simulations presented here, *D*_ZsGreen_=90 *µm*^2^.*s*^*−*1^ and [ZsGreen]=25 *µ*M.

### Protocols for simulating neuronal stimulation

In order to investigate the propagation of Ca^2+^ signals from nodes that contact neuronal spines, we have developed 2 different protocols for our simulations, performed in the geometries presented in Fig 2. As nodes were the site of Ca^2+^ signal initiation Arizono et al. (2020) and as most spines contacted nodes rather than shafts, we have simulated neuronal stimulation in nodes. To simulate neuronal stimulation, IP_3_ and Ca^2+^ were infused in the cytosol at stimulation time. IP_3_ infusion reflects the production of IP_3_ by phospholipase C that results from the activation of *G*_*q*_-G-protein-coupled receptors (GPCRs). Ca^2+^ infusion mimics the influx of Ca^2+^ in the cytosol through Ca^2+^ channels at the plasma membrane. The rate of this neuronal activity-induced Ca^2+^ influx, *k*_*Ca*_, varied within a physiological range of values, from 0 to 1000 *s*^*−*1^ Wu et al. (2018); Brazhe et al. (2018). Signals were recorded both in the stimulated node, Node 1, and in the neighboring node, Node 2.

- In the first protocol, 100 IP_3_ molecules were infused in Node 1, at t=*t*_0_=1s, while Ca^2+^ activity was monitored in Node 1 and in the neighboring node, Node 2 (see e.g Fig 4A). Neuronal activity-induced Ca^2+^ influx was mediated by generic Ca^2+^ channels at the plasma membrane. 25 of those Ca^2+^ channels were placed on the plasma membrane of Node 1, corresponding to a similar density to the IP_3_R density on the ER membrane, and were set to an inactive state. At stimulation time t=*t*_0_, Ca^2+^ channels were set to an active state, resulting in an influx of Ca^2+^ within the cytosol at rate *k*_*Ca*_. At t=*t*_0_ + 1, Ca^2+^ channels were set back to their initial inactive state. Simulations were performed in geometries with varying shaft width d_shaft_.
- In the second protocol, we have investigated signal propagation in the node/shaft geometry depending on shaft width d_shaft_ when several nodes were successively stimulated. In “5nodes” geometries, 50 IP_3_ molecules were infused at *t*_0_=5s, *t*_0_ + *τ*_IP3_, *t*_0_ + 2*τ*_IP3_, *t*_0_ + 3*τ*_IP3_ in Nodes 1, 2, 3 and 4, respectively. During the whole simulation time, Ca^2+^ activity was recorded in Node 5 (see Fig 6). In a subset of simulations, stimulation of Nodes 2, 3 and 4 occurred with a probability 1 − *p*_fail_, with *p*_fail_ ∈ [0, 1].

### Code accessibility

The simulation code, implemented with STEPS 3.5.0, and the meshes are available on ModelDB McDougal et al. (2017) at http://modeldb.yale.edu/266928, access code: lto42@tpk3D?. The original model from Denizot et al. Denizot et al. (2019) is available at http://modeldb.yale.edu/247694.

**Figure 4:**
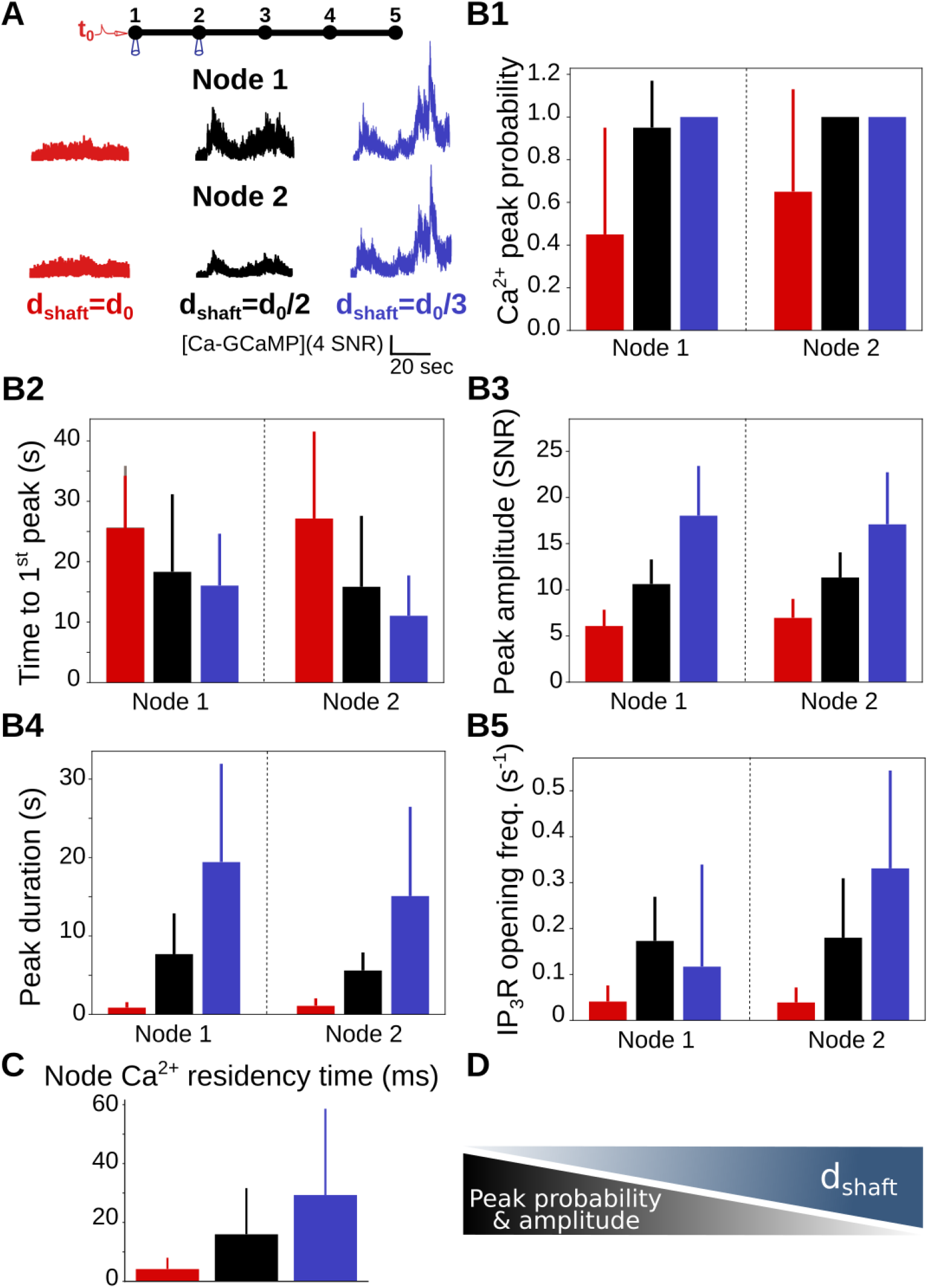
Ca^2+^ peak probability, amplitude and duration increase when shaft width decreases. (*A*) (*Top*) Neuronal stimulation protocol simulated for each geometry: node 1 was stimulated at t=*t*_0_=1s, while Ca^2+^ activity was monitored in node 2. Representative Ca^2+^ traces for shaft width 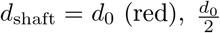 (black) and 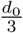 (blue), expressed as SNR (see Methods). (*B*) Quantification of the effect of d_shaft_ on Ca^2+^ signal characteristics Data are represented as mean ± STD, n=20. Ca^2+^ peak probability increases (***, *B1*), Time to 1^*st*^ peak decreases (***, *B2*), peak amplitude (***, *B3*) and duration (***, *B4*) increase when d_shaft_ decreases. (*C*) Ca^2+^ residency time in node 1 increases when d_shaft_ decreases (***, n=300). (*D*) Schematic summarizing the main result from this figure: Ca^2+^ peak probability and amplitude increase when shaft width decreases. The effect of d_shaft_ on each Ca^2+^ signal characteristic was tested using one-way ANOVA. Significance is assigned by * for *p* ≤ 0.05, ** for *p* ≤ 0.01, *** for *p* ≤ 0.001.

### Peak detection and analysis

The same strategy as developed by Denizot et al. Denizot et al. (2019) was used for detecting and analysing Ca^2+^ signals. Briefly, basal concentration of Ca^2+^, [*Ca*]_b_, was defined based on a histogram of the number of Ca^2+^ ions in the absence of neuronal stimulation. Peak initiation corresponded to the time when [Ca^2+^] was higher than the following threshold: [*Ca*]_b_ + *nσ*_Ca_, where *σ*_Ca_ is the standard deviation of [Ca^2+^] histogram in the absence of neuronal stimulation. The value of *n* was set by hand depending on signal/noise ratio of the simulation of interest. Peak termination corresponded to the time when [Ca^2+^] decreased below the peak threshold.

Several parameters were analyzed to characterize Ca^2+^ signals. Peak amplitude, A, corresponds to the maximum [Ca^2+^] measured during the peak duration. It is expressed as signal to noise ratio 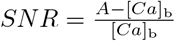. Peak duration corresponds to the time between peak initiation and peak termination. Time to 1^st^ peak corresponds to the delay between the beginning of the simulation and the first peak detection, measured in the cellular compartment of interest. Peak probability corresponds to the fraction of simulations in which at least one peak was detected during simulation time in the region of interest. Ca^2+^ residency time was measured by performing n=300 simulations for each value of d_shaft_, in which only 1 Ca^2+^ ion was added to node 1, without other molecular species. Ca^2+^ residency time corresponds to the time taken for the ion to diffuse away from node 1.

### Organotypic hippocampal slice cultures

All experiments were performed as described in Arizono et al. (2020) and were in accordance with the European Union and CNRS UMR5297 institutional guidelines for the care and use of laboratory animals (Council directive 2010/63/EU). Organotypic hippocampal slices (Gähwiler type) were dissected from 5–7-d-old wild-type mice and cultured 5–8 week in a roller drum at 35°C, as previously described Gähwiler (1981).

### Viral infection

AAV9-GFAP-GCaMP6s Stobart, Ferrari, Barrett, Stobart, et al. (2018) was injected by brief pressure pulses (40ms; 15 psi) into the stratum radiatum of 2-3-week old slices from Thy1-YFP-H (JAX:003782) mice 4-6 weeks prior to the experiment.

### Image acquisition

For Ca^2+^ imaging, we used a custom-built setup based on an inverted microscope body (Leica DMI6000), as previously described in Tonnesen et al. (2011). We used a 1.3 NA glycerol immersion objective equipped with a correction collar to reduce spherical aberrations and thereby allow imaging deeper inside brain tissue Urban et al. (2011). The excitation light was provided by a pulsed diode laser (l = 485 nm, PicoQuant, Berlin, Germany). The fluorescence signal was confocally detected by an avalanche photodiode (APD; SPCM-AQRH-14-FC; PerkinElmer). The spatial resolution of the setup was around 200 nm (in x-y) and 600 nm (z). Confocal time-lapse imaging (12.5 × 25 *µ*m, pixel size 100 nm) was performed at 2Hz for 2.5 min in artificial cerebrospinal fluid containing 125 mM NaCl, 2.5 mM KCl, 1.3 mM MgCl_2_, 2 mM CaCl_2_, 26 mM NaHCO_3_, 1.25 mM NaH_2_PO_4_, 20 mM D-glucose, 1 mM Trolox; 300 mOsm; pH 7.4. Perfusion rate was 2 mL/min and the temperature 32 °C. Hypo-osmotic stress (300 mOsm to 200 mOsm) was applied by perfusing ACSF with reduced NaCl concentration (119 to 69 mM NaCl).

### Experimental Design and Statistical Analysis

For each parameter set, 20 simulations, with different seeds, were generated. Each parameter describing Ca^2+^ dynamics was expressed as mean ± standard deviation. The effect of d_shaft_ on each Ca^2+^ signal characteristic was tested using one-way ANOVA. Comparison between two different conditions was performed using unpaired Student T-test if values followed a Gaussian distribution, Mann-Whitney test otherwise. Significance is assigned by * for *p* ≤ 0.05, ** for *p* ≤ 0.01, *** for *p* ≤ 0.001.

## Results

### Geometrical representation of typical astrocyte processes

In order to investigate the role of the nano-morphology of astrocytic processes on the spatio-temporal properties of Ca^2+^ microdomains, we have designed geometries of typical astrocyte processes, derived from our recent characterization of their ultrastructure at a high spatial resolution (50 nm in x-y) in organotypic hippocampal culture as well as in acute slices and *in vivo* Arizono et al. (2020) (Fig 2A). Geometries consist of alternations of bulbous structures, nodes, connected to each other via cylindrical structures, referred to as shafts. To reproduce the morphological changes of processes associated with cell swelling reported in hypo-osmotic conditions (Fig 3 in Arizono, Inavalli, et al. (2021)), geometries with different shaft width d_shaft_ and constant node width were designed (Fig 2B). In accordance with node-shaft structures observed experimentally Arizono et al. (2020), node width was set to 380 nm. To model astrocytic Ca^2+^ activity with a high spatial resolution while taking into account the randomness of reactions in small volumes, we used the stochastic voxel-based model from Denizot et al. Denizot et al. (2019). The reactions included in the model are presented in Fig 2C and in the Methods section. As the majority of Ca^2+^ signals in astrocytes result from the opening of Inositol 3-Phosphate receptors (IP_3_Rs), located at the membrane of the endoplasmic reticulum (ER) Srinivasan et al. (2015), signals in the model result from the opening of IP_3_Rs (see Denizot et al. Denizot et al. (2019) for discussion on the model’s assumptions and limitations).

### Thin shafts favor node compartmentalization

In order to test whether the geometries designed in this study are a good approximation of the ultrastructure of the gliapil, we have compared molecular diffusion flux in those geometries with those reported experimentally. To do so, we simulated photobleaching experiments and compared our results to experimental results from Arizono et al. Arizono et al. (2020). The principle of bleaching simulations is presented in the Methods section and in Movie S1. Here, we refer to an increased node compartmentalization when the time to recovery after bleaching, *τ*, increases (see Fig 3B).

Bleaching traces in simulations are both qualitatively (Fig 3B) and quantitatively (Fig 3C) similar to experimental bleaching traces, for shaft width *d*_shaft_ = *d*_0_ and 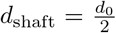. Indeed, no significant difference of *I*_0_ (Fig 3C1), *I*_*inf*_ (Fig 3C2) and *τ* (Fig 3C3) was observed between simulations and experimental traces. Simulations were also performed with 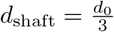. Our simulations successfully reproduce experimental bleaching experiments and suggest that *τ*, and thus node compartmentalization, increases when shaft width decreases (Fig 3C3). This result is not surprising as a decreased shaft width results in a smaller size of the exit point for diffusing molecules from the node. This is similar to e.g dendritic spines, which compartmentalization is increased for thinner spine necks Santamaria et al. (2011); Tonnesen et al. (2014). The geometries that we have designed can thus be considered as a reasonable approximation of the ultrastructure of the gliapil observed experimentally in live tissue.

### Thin shafts enhance Ca^2+^ activity in nodes

80% of astrocyte Ca^2+^ activity occurs in the gliapil Bindocci et al. (2017), which suggests that most neuron-astrocyte communication occurs at fine astrocytic processes. As we observed that a decreased shaft width is associated with a decreased diffusion flux, i.e an increased compartmentalization of nodes, we have tested whether this effect influences Ca^2+^ activity upon neuronal stimulation. To do so, we have first analyzed Ca^2+^ signals resulting from a single neuronal stimulation, which was simulated as an infusion of IP_3_ and Ca^2+^ in the stimulated node, node 1 (see Methods). Those parameters encompass the IP_3_ production by phospholipase C following the activation of *G*_*q*_-G-protein-coupled receptors (GPCRs) resulting from the binding of neuronal glutamate, ATP and noradrenaline to *G*_*q*_ proteins, and Ca^2+^ entry at the plasma membrane through Ca^2+^ channels, ionotropic receptors or the sodium/calcium exchanger (NCX) functioning in reverse mode Ahmadpour et al. (2021); Semyanov et al. (2020). Signals were recorded both in the stimulated node, node 1, and in the neighboring node, node 2 (Movie S2). Representative Ca^2+^ traces in node 2 for 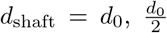 and 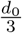 are displayed in Fig 4A. Our first result is that Ca^2+^ peak probability increases when d_shaft_ decreases (Fig 4B1). The time to 1^*st*^ peak increases with d_shaft_ (Fig 4B2). By contrast, peak amplitude (Fig 4B3) and duration (Fig 4B4) increase when d_shaft_ decreases. To better understand the mechanisms responsible for the increased Ca^2+^ peak probability, amplitude and duration when d_shaft_ decreases, we measured the frequency of *IP*_3_*R* opening in nodes 1 and 2. The frequency of *IP*_3_*R* opening increases when d_shaft_ decreases (Fig 4B5). Note that the duration of *IP*_3_*R* opening and the number of *IP*_3_*R*s open per Ca^2+^ peak did not vary with d_shaft_. The increased *IP*_3_*R* opening frequency associated with small values of d_shaft_ probably results from the increased residency time of molecules in nodes connected to thin shafts (Fig 4C). Indeed, a thin shaft can “trap” Ca^2+^ and *IP*_3_ longer in the node, thus locally increasing the probability of *IP*_3_*R*s to open, resulting in larger Ca^2+^ peaks. For more details, the reader can refer to the theoretical work investigating the narrow escape problem for diffusion in microdomains Schuss et al. (2007). Nodes connected to thinner shafts, despite being characterized by a lower diffusion flux (Fig 3), could thus consist of signal amplification units, favoring the generation of larger signals, therefore increasing Ca^2+^ peak probability, amplitude and duration both in the stimulated and in neighboring nodes.

To identify the cause of the increased Ca^2+^ activity when shaft width decreases (Fig 4), we performed simulations in which we altered the stimulated node (Figure 3 S2D-E), Ca^2+^ influx at the plasma membrane (Figure S3) and boundary conditions (Figure S4). Those parameters did not affect the observed effects of d_shaft_ on Ca^2+^ dynamics. Furthermore, we repeated simulations of Fig 4 with constant cytosolic volume and constant number of IP_3_Rs, irrespective of the value of d_shaft_ (Figure S5). Our results highlight that the relevant parameter responsible for the observed effects of d_shaft_ on Ca^2+^ signal characteristics is the node/shaft width ratio or the cytosolic volume rather than d_shaft_ itself. Spontaneous Ca^2+^ signals were affected by shaft width in the same way as neuronal-induced Ca^2+^ signals (Figure S6) and, in particular, reproduced the increase of the amplitude ratio of spontaneous Ca^2+^ signals between node 2 and node 1 with shaft width observed in hippocampal organotypic cultures Arizono et al. (2020). Note that ER morphology, in particular ER surface area (Figure S7, S8 and S9), and Ca^2+^ buffering by Ca^2+^ indicators (Figure S10), consistent with previous reports Denizot et al. (2019); Bartol et al. (2015); Majewska et al. (2000), also altered local Ca^2+^ activity. Overall, our results suggest that a decreased shaft width, resulting in a decreased diffusion efflux from nodes, increases Ca^2+^ peak probability, amplitude and duration. Conversely, the swelling of fine processes, resulting in an increase of shaft width, attenuates local Ca^2+^ peak probability, amplitude and duration.

### Ca^2+^ imaging confirms that swelling attenuates local spontaneous Ca^2+^ activity

To validate the role of thin shafts suggested by our model’s predictions, we tried to recreate the widening of shaft width in experimental conditions. Coincidentally, our recent super-resolution study revealed that the nano-architecture of fine processes is remodeled in hypo-osmotic conditions, where shaft width increased while node width remained unaltered (see Fig 3 in Arizono, Inavalli, et al. (2021)). While hypo-osmotic conditions undoubtedly cause many physiological changes to astrocytes, it is the closest experimentally available model to test our model predictions. We thus performed experimental measurements of Ca^2+^ activity in fine astrocytic processes under basal and hypo-osmotic conditions. To record Ca^2+^ signals in fine branchlets, we used confocal microscopy in organotypic cultures, which provide a high level of optical access and sample stability in live tissue (resolution ≈ 200 nm in x-y versus ≈ 500 nm for two-photon microscopy). In accordance with our model’s predictions, Ca^2+^ peak amplitude and duration were lower in hypo-osmotic compared to basal conditions (Fig 5A-D). Such differences were not observed in the absence of HOC (Fig 5E-F). Overall, our experimental results confirm that the complex nano-architecture of fine astrocytic processes and its alteration, such as cell swelling, shapes local Ca^2+^ activity.

**Figure 5:**
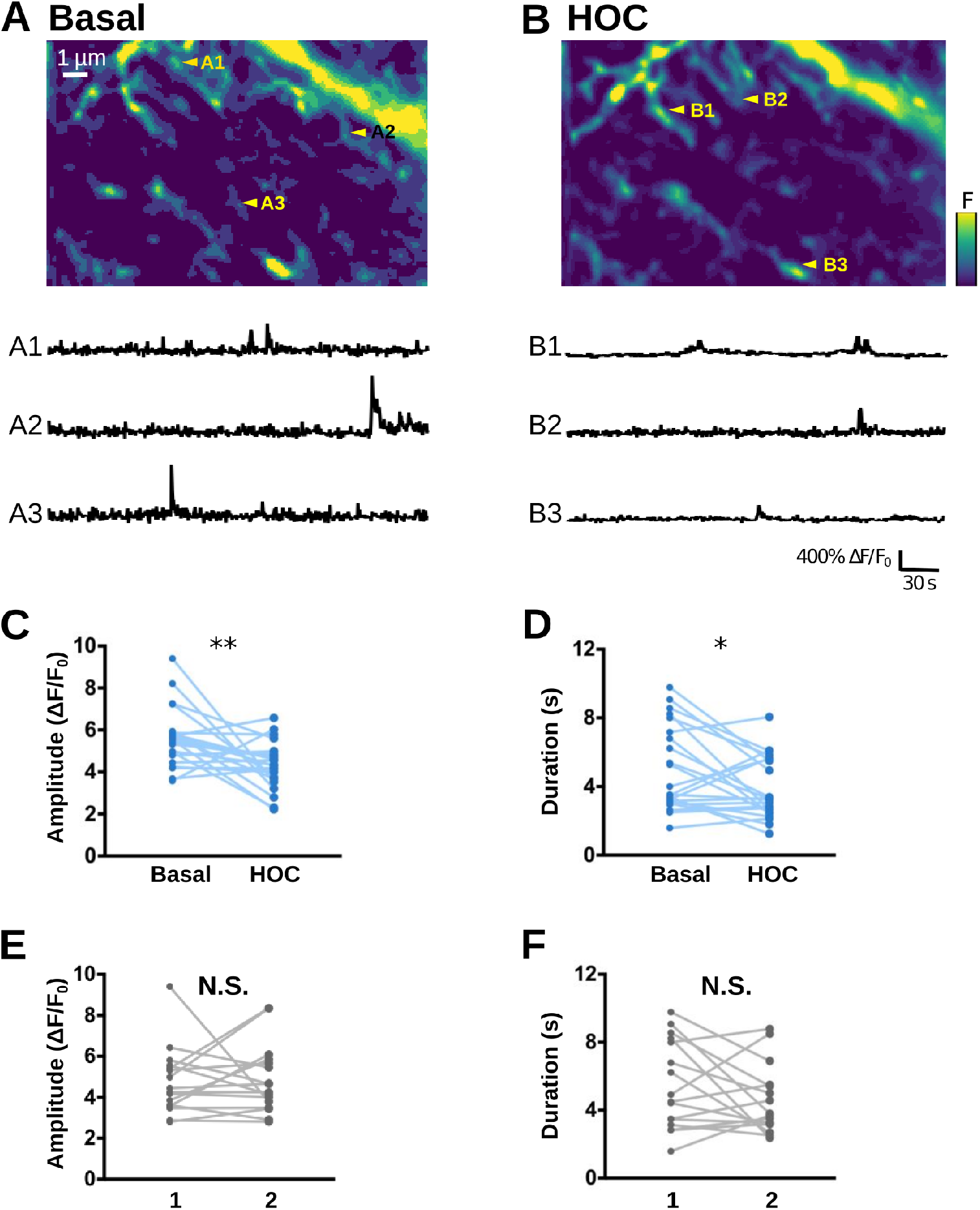
Ca^2+^ imaging confirms that swelling attenuates local spontaneous Ca^2+^ activity. (*A, B*) (Top) Confocal images of the astrocytic spongiform domain expressing GCaMP6s at baseline (basal, A) and in hypo-osmotic condition (HOC, B), measured in organotypic hippocampal cultures (resolution: 200 nm in x-y, 600 nm in z). (Bottom) Representative traces of spontaneous Ca^2+^ events from ROIs indicated in A (A1-A3) and B (B1-B3). (*C, D*) Ca^2+^ peak amplitude (C) and duration (D) of spontaneous Ca^2+^ events are significantly smaller in hypo-osmotic conditions (HOC) compared to basal conditions (Basal). (*E, F*) Amplitude (E) and duration (F) of spontaneous Ca^2+^ events do not significantly vary when measured twice in a row (1, 2) in the absence of HOC. Lines represent measurements in the same cell, before and after applying hypo-osmotic stress. Significance is assigned by * for *p* ≤ 0.05, ** for *p* ≤ 0.01, *** for *p* ≤ 0.001.

### Thin shafts favor more robust signal propagation

As a single branchlet communicates with multiple dendritic spines, which can function independently or belong to a cluster of co-active synapses Arizono et al. (2020); Reichenbach et al. (2010); Witcher et al. (2007); Cali et al. (2019); Semyanov et al. (2020), the frequency of node stimulation within the branchlet can vary drastically. Thus, we have tested how node stimulation frequency affects Ca^2+^ activity in the branchlet. To do so, we have performed simulations in which neighboring nodes were repeatedly stimulated after a time period *τ*_IP3_, that varied from 50 ms to 5 s, while Ca^2+^ signals were recorded in a remote node, node 5 (Fig 6A). Neuronal stimulation is simulated as an infusion of 50 IP_3_ molecules in the stimulated node. Representative Ca^2+^ traces in node 5 in branchlets with various shaft widths d_shaft_ are presented in Fig 6A for *τ*_IP3_=250 and 3000 ms. Our first notable result is that the time to 1^*st*^ peak in node 5 decreases with d_shaft_, whatever the value of *τ*_IP3_ (Fig 6B1). More specifically, time to 1^*st*^ peak is higher for d_shaft_ =*d*_0_ compared to both 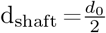 and 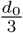, while differences between 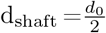 and 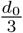 are not as striking. Moreover, the difference between 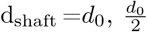 and 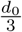 increases with *τ*_IP3_. This suggests that geometries with d_shaft_ =*d*_0_ better discriminate slow from fast frequency of node stimulation compared to geometries with thinner shafts. Geometries with d_shaft_ =*d*_0_ are further characterized by a lower Ca^2+^ peak probability in node 5 compared to geometries with 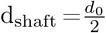 and 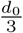 (Fig 6B2). More precisely, Ca^2+^ peak probability decreases as *τ*_IP3_ increases for d_shaft_ =*d*_0_, which was observed independently of our boundary conditions (Figure S11, see Methods). This suggests that geometries with larger shafts could be associated with decreased signal propagation to remote nodes in case of repeated node stimulation at low frequency (*τ*_IP3_ *>* 2 s).

**Figure 6:**
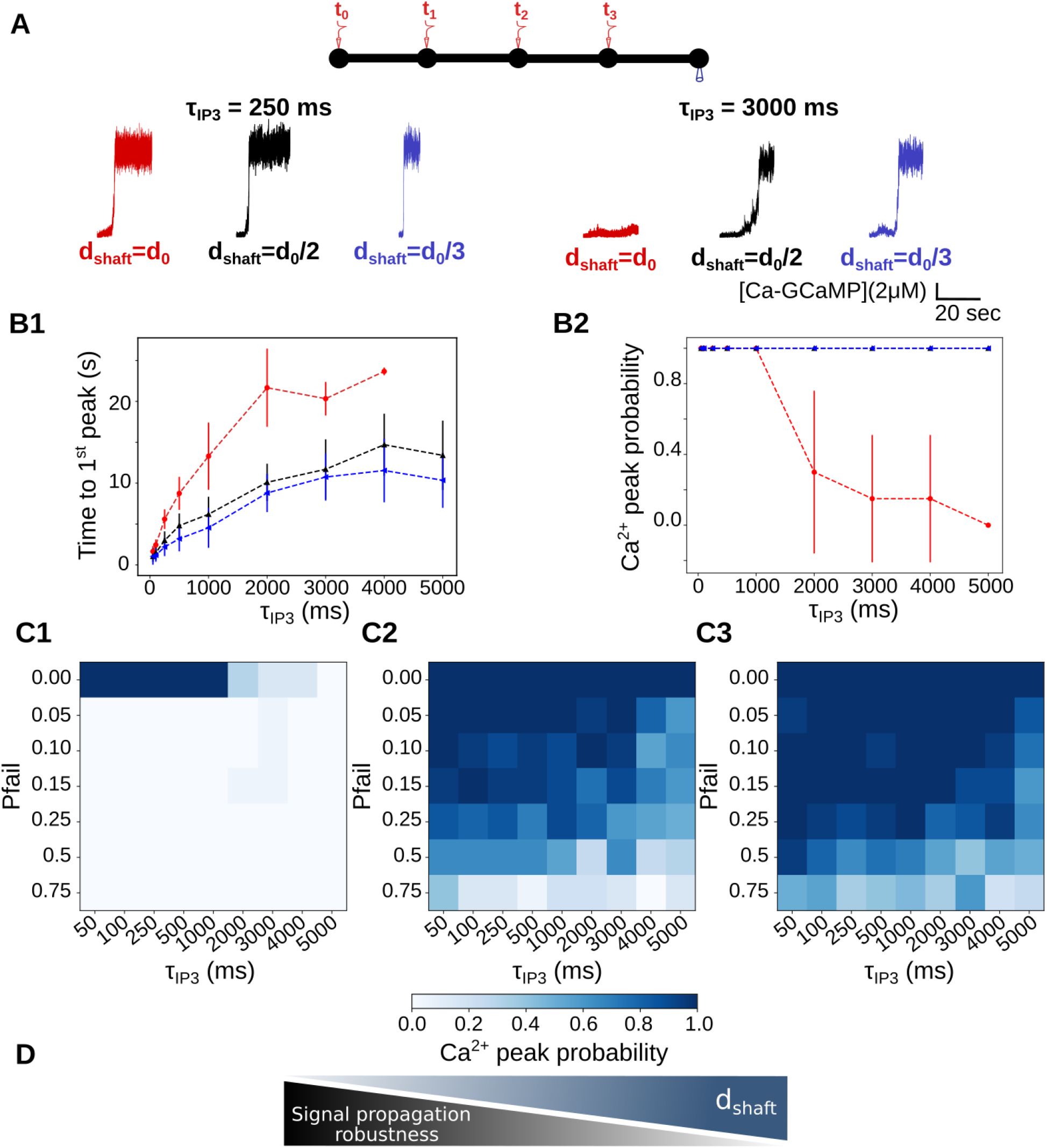
Thin shafts favor a more robust signal propagation upon repeated neurotransmitter release events. (*A*) (*Top*) Neuronal stimulation protocol: node 1 is stimulated at t=*t*_0_=5s, node 2 at *t*_0_ + *τ*_IP3_, node 3 at *t*_0_ + 2*τ*_IP3_ and node 4 at *t*_0_ + 3*τ*_IP3_, *k*_*Ca*_=0 *s*^*−*1^. Ca^2+^ activity is recorded in node 5. (*Bottom*) Representative Ca^2+^ traces in node 5 for shaft width *d*_shaft_ = *d*_0_ (red), 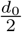 black) and 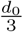 (blue), with *τ*_IP3_=250 ms (left) and 3000 ms (right), expressed as SNR (see Methods). (*B1*) Time to 1^*st*^ peak increases with *τ*_IP3_ for 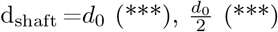 and 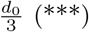. T-tests revealed that for any value of *τ*_IP3_, time to 1^*st*^ peak is higher for d_shaft_ =*d*_0_ compared to 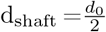 and 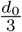. Time to 1^*st*^ peak is significantly higher when 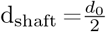 compared to 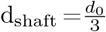, for most values of *τ*_IP3_ (p=0.032 *, 0.0025 **, 0.034 *, 0.016 * and 0.019 * for *τ*_IP3_=250, 500, 1000, 4000 and 5000 ms, respectively). (*B2*) Ca^2+^ peak probability in node 5 is lower for d_shaft_ =*d*_0_ compared to 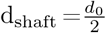 and 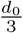. Ca^2+^ peak probability decreases as *τ*_IP3_ increases for d_shaft_ =*d*_0_ (***). (*C*) Ca^2+^ peak probability in node 5 (colorbar) as a function of *τ*_IP3_ and of the probability of failure of node stimulation *p*_fail_, for d_shaft_ =*d*_0_ (C1), 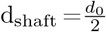 (C2) and 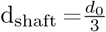 (C3), with *p*_fail_ ∈ [0, 1]. (*D*) Schematic summarizing the main conclusion of this figure: decreased shaft width allows signal propagation despite omitted node stimulation, thus favoring more robust signal propagation. Data are represented as mean ± STD, n=20 for each value of d_shaft_ and of *τ*_IP3_. Lines in panel B are guides for the eyes. The effect of d_shaft_ on each Ca^2+^ signal characteristic was tested using one-way ANOVA. Significance is assigned by * for *p* ≤ 0.05, ** for *p* ≤ 0.01, *** for *p* ≤ 0.001.

For *τ*_IP3_=4s and 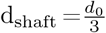, signals were detected in node 5 11.55 ± 3.89 s after the stimulation of node 1, which means that they occurred before the stimulation of node 4 (t=*t*_0_ + 12s for *τ*_IP3_=4s). This phenomenon was not observed for d_shaft_ =*d*_0_, for which time to 1^st^ peak when *τ*_IP3_=4s was 23.67 ± 0.47 s. This suggests that for 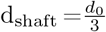, contrary to d_shaft_ =*d*_0_, one node stimulation could be omitted without having any consequence on Ca^2+^ peak probability in node 5. In order to test this hypothesis, we have performed simulations in which the stimulation of nodes 2, 3 and 4 occurred with a given probability of failure *p*_fail_. Simulations were performed for *p*_fail_=0, 0.05, 0.1, 0.15, 0.25 and 0.75. Ca^2+^ peak probability in node 5, depending on *p*_fail_ and on *τ*_IP3_ is presented in Fig 6C, for d_shaft_ =*d*_0_ (Fig 6C1), 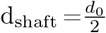 (Fig 6C2) and 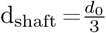 (Fig 6C3). As expected, Ca^2+^ peak probability, despite high values of *p*_fail_, increases when d_shaft_ decreases. Thus, thin shafts can favor signal propagation by allowing the omission of a node stimulation. In that sense, geometries displaying thin shafts are characterized by a more robust signal propagation (Fig 6D).

Together, our results suggest that, in the context of repeated node stimulation, thin shafts are associated with an increase of Ca^2+^ peak probability in more remote nodes, with an earlier signal onset, suggesting increased signal propagation. Astrocytic processes with thicker shafts (here d_shaft_ =*d*_0_), such as observed in hypo-osmotic conditions Arizono, Inavalli, et al. (2021), are associated with lower signal propagation in case of low stimulation frequency (time period *>* 2 s), potentially favoring the formation of local Ca^2+^ hotspots. Our results suggest that geometries with thick shafts could impair signal propagation when a branchlet is stimulated at a low frequency. In that sense, astrocyte branchlets with thicker shafts would be better detectors of the surrounding level of neuronal activity. By contrast, branchlets with thin shafts would be less discriminating and provide more robust signal propagation.

## Discussion

Fine astrocytic processes are responsible for most astrocytic Ca^2+^ signals Bindocci et al. (2017) and are preferential sites of neuron-astrocyte communication Arizono et al. (2020). A better understanding of the mechanistic link between their morphology and the spatio-temporal properties of local Ca^2+^ signals is crucial, yet hard to test experimentally. Here, we perform reaction-diffusion simulations in idealized morphologies of astrocytic processes derived from 3D super-resolution microscopy to investigate the effect of astrocyte nanoscale morphology on Ca^2+^ activity in the gliapil. Our simulation results indicate that the nanoscale morphological features of astrocytic processes effectively increase the peak probability, duration, amplitude and propagation of Ca^2+^ signals. Conversely, the alteration of the node-shaft arrangement of the spongiform domain associated with astrocyte swelling attenuates local Ca^2+^ activity and signal propagation (Fig **??**). Our simulation results, in accordance with experimental data, suggest that thin shafts effectively decrease diffusion flux, resulting in an increased compartmentalization of biochemical signals in nodes. Thus, nodes, similarly to dendritic spines Santamaria et al. (2011), act as diffusion traps when shaft width is low. Note that, more than the value of shaft width itself, our results emphasize the effect of the ratio between node and shaft diameter on Ca^2+^ activity. The simple geometries that we have evaluated in this study could be used to build a more comprehensive model of the spongiform structure to simulate Ca^2+^ activity in the entire astrocyte. By recording the molecular interactions resulting in Ca^2+^ signals upon neuronal stimulation in small cellular compartments of the gliapil, which cannot be performed experimentally, our simulation results shed light on the mechanisms by which the nano-architecture of astrocytic processes influences the frequency, amplitude and propagation of local Ca^2+^ signals at tripartite synapses in health and disease.

Experimental Ca^2+^ recordings of astrocyte activity have established that astrocyte processes display both highly localized microdomain signals and propagating Ca^2+^ waves Srinivasan et al. (2015); Bindocci et al. (2017). Our simulations suggest that the morphology of the cell and of its organelles can strongly influence the formation of these patterns of astrocytic Ca^2+^ signaling. Notably, thinner shafts allow less discriminating and more robust signal propagation upon repeated stimuli compared to larger shafts. On the contrary, geometries with thick shafts seem to be more discriminating, potentially favoring the propagation of signals resulting from repeated stimuli from co-active synapses. Cellular morphology thus emerges as a key parameter that regulates the active propagation of Ca^2+^ signals. The ultrastructure of the spongiform domain of astrocytes is very complex, characterized by abundant branching points, conferring a reticular morphology Arizono et al. (2020). Those branching points are reportedly sometimes arranged into ring-like structures, although their occurrence and shape are still debated and could differ depending on the brain region under study Arizono et al. (2020); Panatier et al. (2014); Kiyoshi et al. (2020); Salmon et al. (2021); Arizono & Nägerl (2021). The effect of this reticular ultrastructure on the propagation of Ca^2+^ signals remains to be uncovered. Further characterization of the shape of fine astrocytic processes of the spongiform domain, their variability as well as their connectivity to the neighboring synapses are thus required. Pairing those observations with biophysically-detailed models such as the one presented in this study stands to deepen our understanding of the roles of astrocytic and neuronal morphology at tripartite synapses on neuron-astrocyte communication.

In neurons, both experimental Yuste et al. (2000); Noguchi et al. (2005); Tonnesen et al. (2014) and modelling Schmidt & Eilers (2009); Biess et al. (2007); Simon et al. (2014); Bell et al. (2019); Holcman & Schuss (2005, 2011); Santamaria et al. (2011); Cugno et al. (2019) studies have suggested that thin spine necks favor the compartmentalization of Ca^2+^ signals within the spine head. This compartmentalization of synapses allows neurons to discriminate various inputs and to process information locally Wybo et al. (2019); Poirazi & Papoutsi (2020), increasing the computational power of the neuronal circuits. According to our simulation results, nodes connected to thin shafts could favor the emergence of large signals at the site of neuron-astrocyte communication. Interestingly, we further propose that those amplified signals in nodes, instead of resulting in Ca^2+^ hotspots, favor active signal propagation. Fine astrocytic processes encounter morphological rearrangements that are activity-dependent, which notably influence synaptic maturation, efficacy and spine stability Theodosis et al. (2008); Zhou et al. (2019); Henneberger et al. (2020). Our study sheds light on the influence of rearrangements of the reticular morphology of fine processes on signal computation by astrocytes. Further investigation manipulating astrocyte morphology *in situ* as well as *in vivo* is required to better characterize the variability of astrocyte ultrastructure and the associated integration of Ca^2+^ signals.

The morphology of the complex spongiform domain of astrocytes is highly dynamic, subject to activity-dependent as well as pathological remodeling. Our study, providing mechanisms by which an altered astrocyte morphology influences neuron-astrocyte communication at the nanoscale, gives new insights into the involvement of astrocytes in brain function in health and disease.

## Acknowledgements

This work was funded by the Okinawa Institute of Science and Technology Graduate University and by JSPS (Japan Society for the Promotion of Science) Postdoctoral Fellowship for Research in Japan (Standard, P21733). We thank Iain Hepburn and Weiliang Chen of the Computational Neuroscience Unit, OIST, Japan for discussion on 3D meshes and STEPS.

