## Supplementary Information for "Control of Ca^2+^ signals by astrocyte nanoscale morphology at tripartite synapses"

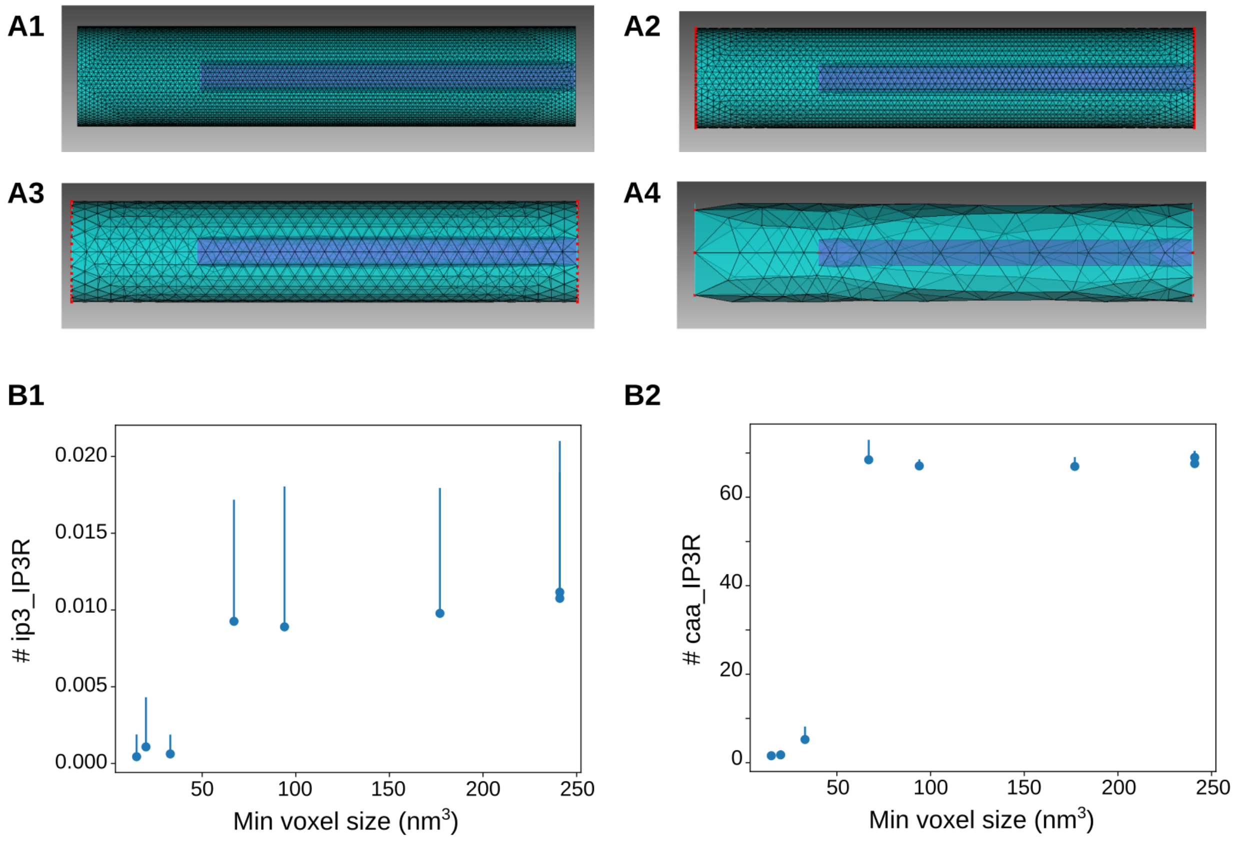


**Figure S1. Sensitivity study of the effect of voxel size on** $Ca^{2+}$ **signals.** (*A*) Simulations were performed in a cylinder geometry with various numbers of tetrahedra. 4 examples are presented here, "$Cyl_1$" (A1), "$Cyl_3$" (A2), "$Cyl_5$" (A3) and "$Cyl_8$" (A4). Please refer to Table **??** for details on the characteristics of each mesh. (B) Study of the average number of IP3R in the ip3_IP3R state ($IP_3$ bound to its binding site on $IP_3R$, B1) and in the caa_IP3R state ($Ca^{2+}$ bound to the activating site of $IP_3R$, B2) depending on the mimimum voxel size of the mesh.

| Geom | # tet | $V_{\text{mean}}$ | $V_{\text{std}}$ | $V_{\text{max}}$ | $V_{\text{min}}$ | Diff ($nm^3$) | Diff (%) |
|---|---|---|---|---|---|---|---|
| $Cyl_1$ | 372148 | 84 | 25 | 296 | 15 | 281 | 335 |
| $Cyl_2$ | 274233 | 115 | 34 | 389 | 20 | 369 | 321 |
| $Cyl_3$ | 125927 | 249 | 71 | 761 | 33 | 728 | 292 |
| $Cyl_4$ | 40077 | 782 | 229 | 2200 | 67 | 2133 | 273 |
| $Cyl_5$ | 11523 | 2700 | 790 | 7521 | 177 | 7344 | 272 |
| $Cyl_6$ | 3103 | 9900 | 3200 | 24000 | 94 | 23906 | 241 |
| $Cyl_7$ | 1417 | 21000 | 11000 | 62000 | 241 | 61759 | 294 |
| $Cyl_8$ | 945 | 32000 | 28000 | 145000 | 241 | 144759 | 452 |

**Table S1. Characteristics of meshes used to investigate the effect of voxel size on** $Ca^{2+}$ **signals.** Cylinder geometries were generated with various total number of tetrahedra (geometry $Cyl_{\text{i}}$ with $i \in [1, 8]$, see also Fig S1A), resulting in different voxel sizes. $V_{\text{mean}}$ corresponds to the average voxel volume, $V_{\text{std}}$ to its standard deviation, $V_{\text{max}}$ to the maximum voxel volume and $V_{\text{min}}$ to the minimum voxel volume in the mesh. Volumes are expressed in $nm^3$. Diff ($nm^3$) corresponds to $V_{\text{max}} - V_{\text{min}}$ while Diff (%) is the ratio $\frac{V_{\text{max}} - V_{\text{min}}}{V_{\text{mean}}} * 100$

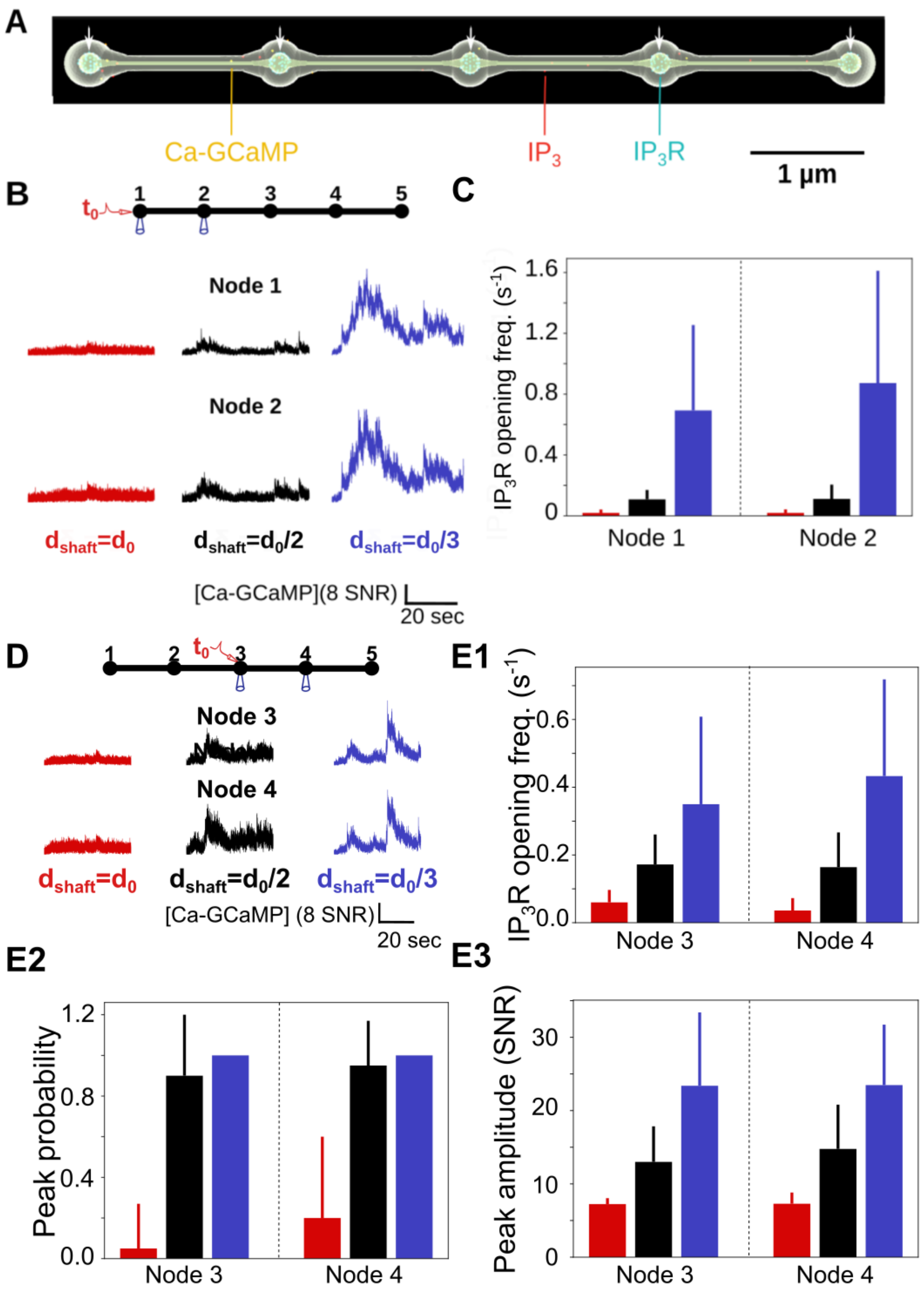


**Figure S2. Sensitivity study of the effect of shaft width on local $Ca^{2+}$ signals upon**

**single node stimulation.** ($A$) Screenshot of a simulation in the "5nodes" $d_{\text{shaft}} = \frac{d_0}{3}$ geometry. In those simulations, $IP_3Rs$ are only positioned on the ER membrane in nodes (white arrows). The number of $IP_3R$ channels is 300 for all values of $d_{\text{shaft}}$. ($B$) (*Top*) Node 1 was stimulated at t=$t_0$=1s, while $Ca^{2+}$ activity was monitored in nodes 1 and 2. (*Bottom*) Representative $Ca^{2+}$ traces in node 1 and node 2 for $d_{\text{shaft}}$ =$d_0$ (red), $\frac{d_0}{2}$ (black) and $\frac{d_0}{3}$ (blue). ($C$) The frequency of $IP_3R$ opening increases with $d_{\text{shaft}}$ decreases in nodes 1 (***) and 2 (***). ($D$) (*Top*) In a second set of simulations, node 3 was stimulated at t=$t_0$=1s ($k_{Ca}$=0 $s^{-1}$), while $Ca^{2+}$ activity was monitored in nodes 3 and 4. (*Bottom*) Representative [Ca-GCaMP] traces in nodes 3 and 4, expressed as SNR (see Methods), for $d_{\text{shaft}}$ =$d_0$ (red), $\frac{d_0}{2}$ (black) and $\frac{d_0}{3}$ (blue). (*E1*) The frequency of $IP_3R$ opening increases when $d_{\text{shaft}}$ decreases in node 3 (***) and 4 (***). (*E2*) $Ca^{2+}$ peak probability increases when $d_{\text{shaft}}$ decreases in node 3 (***) and 4 (***). (*E3*) Peak amplitude increases when $d_{\text{shaft}}$ decreases in node 3 (***) and 4 (***). Data are represented as mean ± STD, n=20 for each geometry. The effect of $d_{\text{shaft}}$ on each $Ca^{2+}$ signal characteristic was tested using one-way ANOVA. Significance is assigned by * for $p \leq 0.05$, ** for $p \leq 0.01$, *** for $p \leq 0.001$.

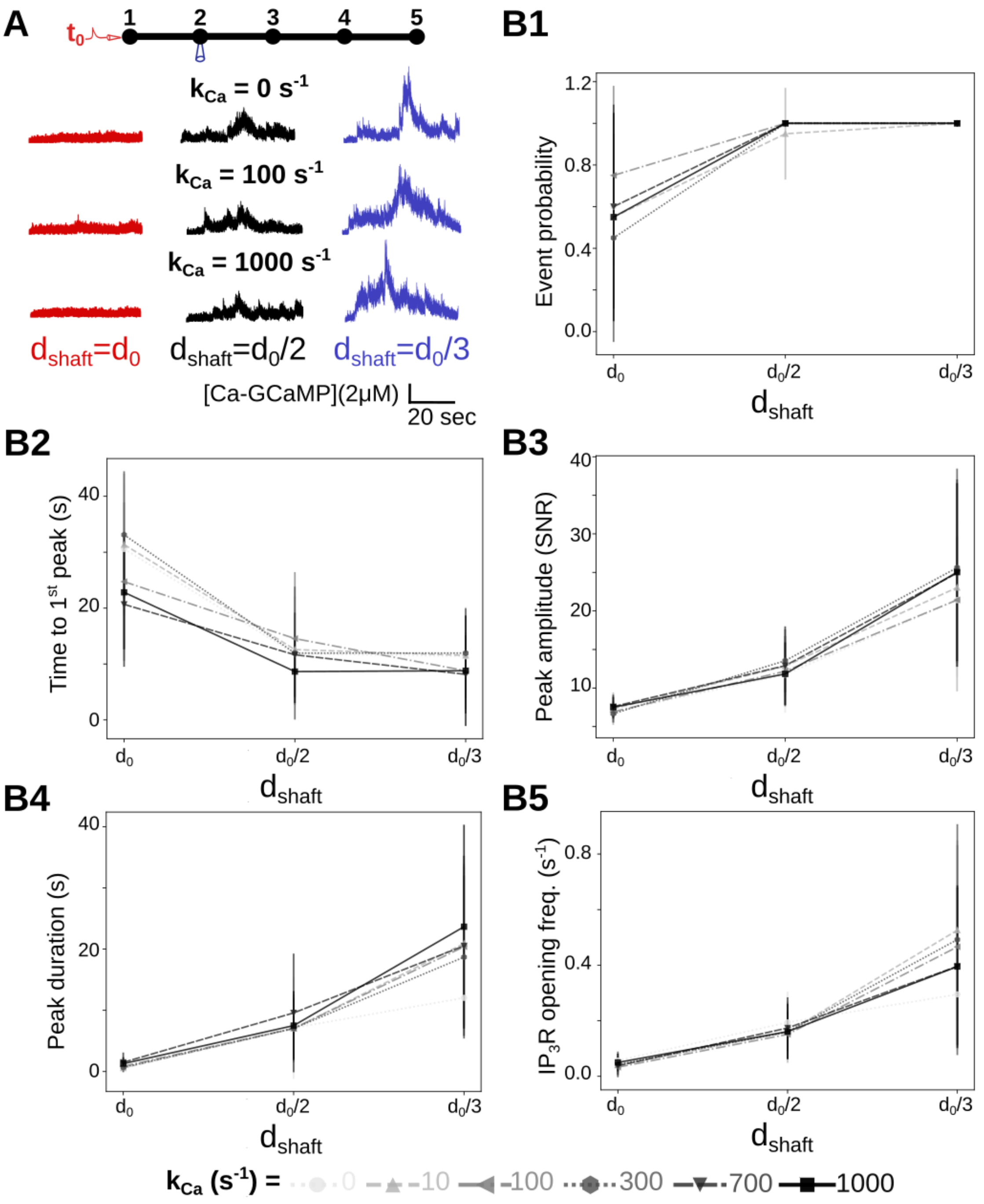


**Figure S3. Effect of $Ca^{2+}$ influx rate at the plasma membrane on local $Ca^{2+}$ signals.** *A*) (*Top*) Neuronal stimulation protocol simulated for each geometry: node 1 was stimulated at t=$t_0$=1s, while $Ca^{2+}$ activity was monitored in node 2. Representative $Ca^{2+}$ traces for shaft width $d_{\text{shaft}} = d_0$ (red), $\frac{d_0}{2}$ (black) and $\frac{d_0}{3}$ (blue), with a $Ca^{2+}$ influx rate at the plasma membrane $k_{Ca}$=0 (top), 100 (middle) and 1000 $s^{-1}$ (bottom), expressed as SNR (see Methods). (*B*) Quantification of the effect of $d_{\text{shaft}}$ on $Ca^{2+}$ signal characteristics for $k_{Ca}$=0, 10, 100, 300, 700 and 1000 $s^{-1}$. Data are represented as mean ± STD, n=20 for each set of parameters tested. $Ca^{2+}$ peak probability increases (***, *B1*), Time to $1^{st}$ peak decreases (***, *B2*), peak amplitude (***, *B3*) and duration (***, *B4*) increase when $d_{shaft}$ decreases, for $k_{Ca}$=0, 10, 100, 300, 700 and 1000 $s^{-1}$. $Ca^{2+}$ peak characteristics did not statistically vary with $k_{Ca}$. The effect of $k_{Ca}$ on each $Ca^{2+}$ signal characteristic was tested using one-way ANOVA.

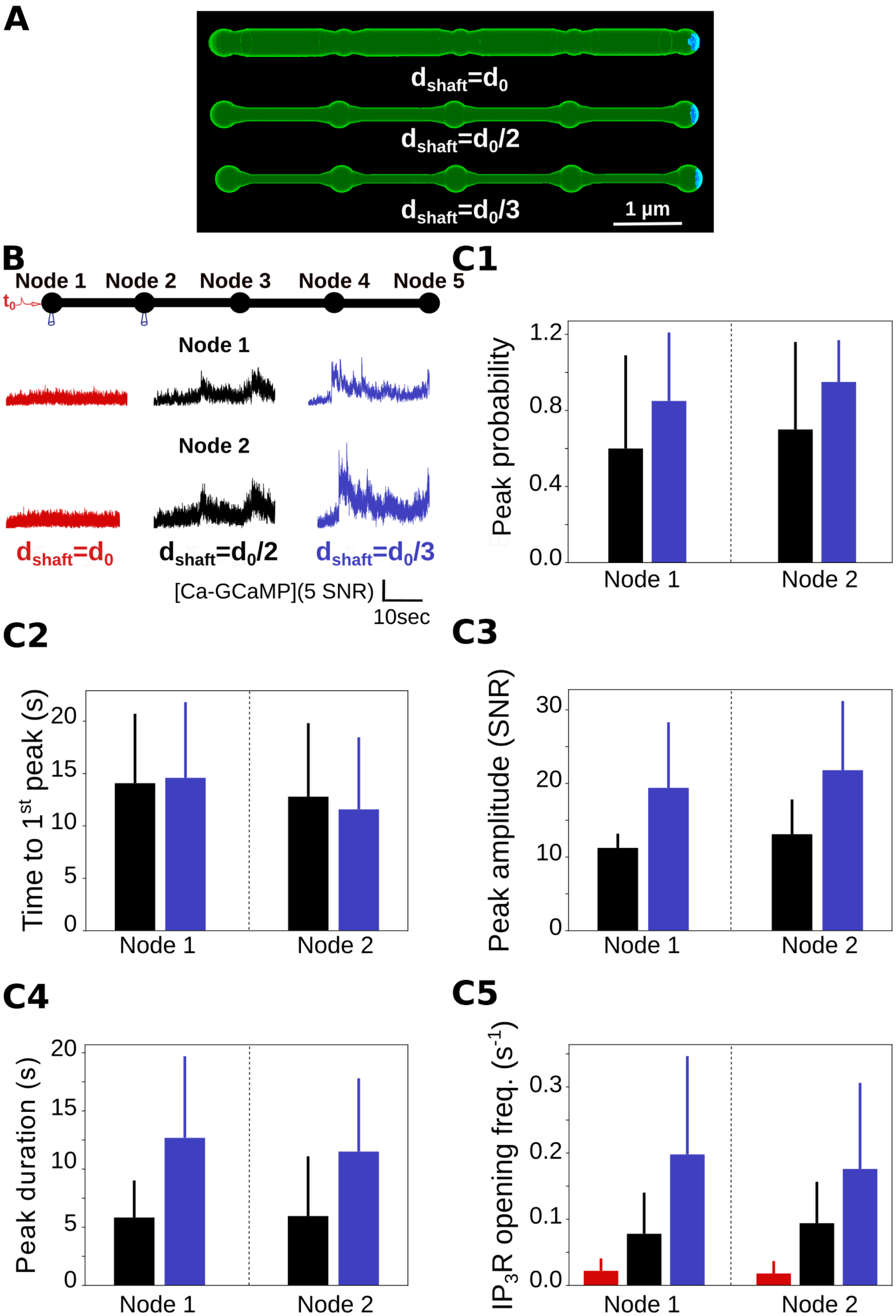

A
$d_{shaft}=d_0$
$d_{shaft}=d_0/2$
$d_{shaft}=d_0/3$
1 μm
B
Node 1
Node 2
Node 3
Node 4
Node 5
$t_0$
Node 1
Node 2
$d_{shaft}=d_0$
$d_{shaft}=d_0/2$
$d_{shaft}=d_0/3$
[Ca-GCaMP](5 SNR)
10sec
C1
Peak probability
1.2
0.8
0.4
0.0
Node 1
Node 2
C2
Time to $1^{st}$ peak (s)
20
15
10
5
0
Node 1
Node 2
C3
Peak amplitude (SNR)
30
20
10
0
Node 1
Node 2
C4
Peak duration (s)
20
15
10
5
0
Node 1
Node 2
C5
$IP_3R$ opening freq. ($s^{-1}$)
0.3
0.2
0.1
0.0
Node 1
Node 2

**Figure S4. Sensitivity study of the effect of boundary conditions on local $Ca^{2+}$ activity upon single node stimulation.** (*A*) Geometries used in the simulations were the "5nodes" geometries (Fig1B). Boundary conditions were absorbing at the extremity of node 5 (blue), while the remaining boundaries were reflective (green). This mimics the connection of the modeled astrocyte branchlet to the rest of the cell. (*B*) (*Top*) Neuronal stimulation protocol simulated for each geometry: node 1 was stimulated at t=$t_0$=1s ($k_{Ca}$=0 $s^{-1}$), while $Ca^{2+}$ activity was monitored in nodes 1 and 2. (*Bottom*) Representative [Ca-GCaMP] traces in nodes 1 and 2, expressed as SNR (see Methods), for $d_{shaft} = d_0$ (red), $\frac{d_0}{2}$ (black) and $\frac{d_0}{3}$ (blue). (*C1*) $Ca^{2+}$ peak probability increases when $d_{shaft}$ decreases in nodes 1 (***) and 2 (***). (*C2*) Time to $1^{st}$ peak does not change with $d_{shaft}$, both in node 1 (p-value=0.85) and 2 (p-value=0.90). (*C3*) Peak amplitude increases when $d_{shaft}$ decreases in nodes 1 (p-value=0.0056 **) and 2 (p-value=0.0021 **). (*C4*) Peak duration increases when $d_{shaft}$ decreases in nodes 1 (p-value=0.0051 **) and 2 (p-value=0.014 *). (*C5*) The frequency of $IP_3R$ opening increases when $d_{shaft}$ decreases in both nodes 1 and 2 (***). Note that there were no $Ca^{2+}$ peak in simulations with $d_{shaft} = d_0$. $Ca^{2+}$ peak characteristics were not statistically different between node 1 and 2. Data are represented as mean ± STD, n=20 for each geometry. The effect of $d_{shaft}$ on each $Ca^{2+}$ signal characteristic was tested using one-way ANOVA. Significance is assigned by * for $p \leq 0.05$, ** for $p \leq 0.01$, *** for $p \leq 0.001$.

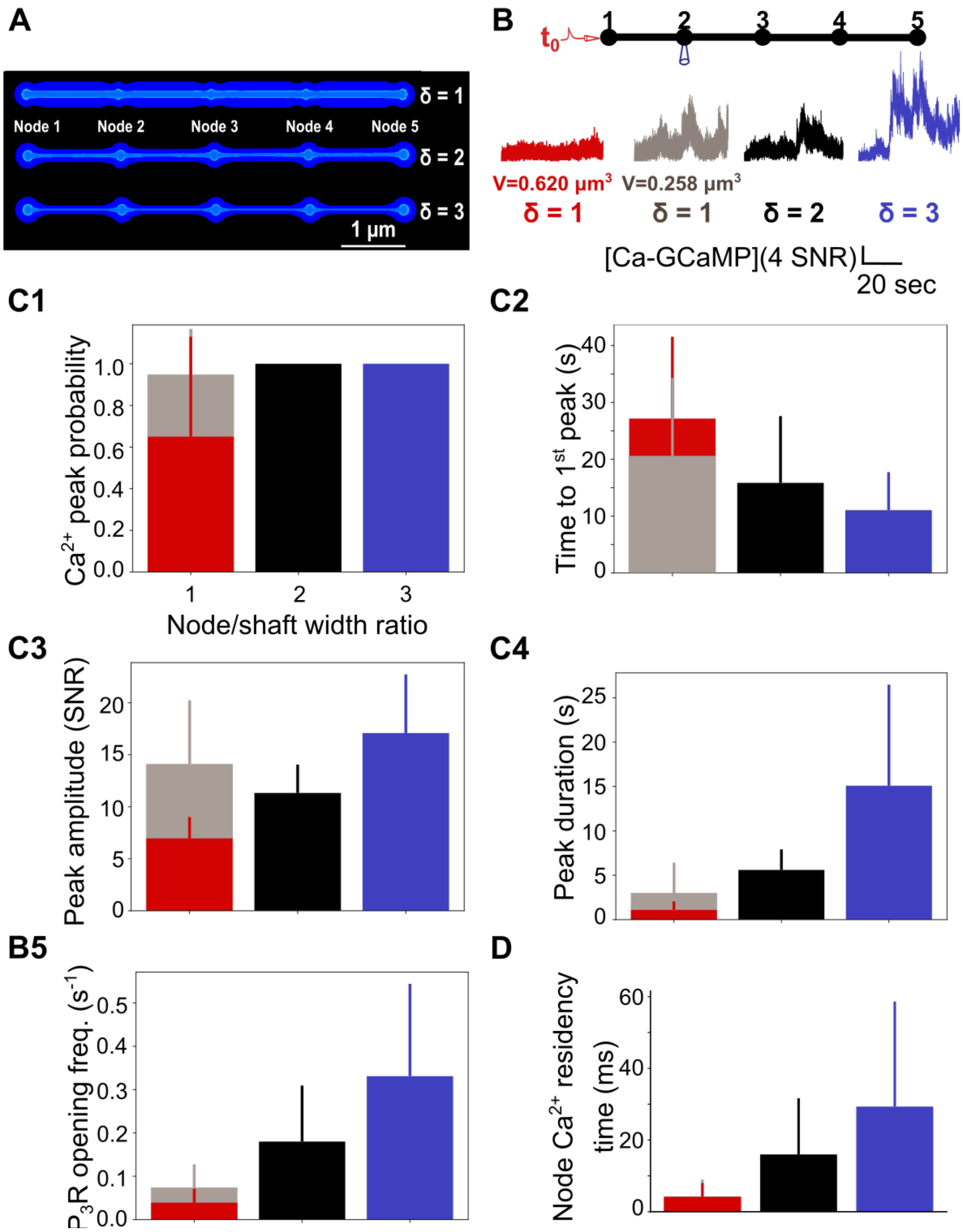


**Figure S5. Node/shaft width ratio and node volume control local** $Ca^{2+}$ **activity.** (*A*) Screenshots from STEPS visualization toolkit revealing meshes used in this study, that contain 5 identical nodes and 4 identical shafts. (*B*) (*Top*) Neuronal stimulation protocol simulated for each geometry: node 1 was stimulated at t=$t_0$=1s ($k_{Ca}$=0 $s^{-1}$), while $Ca^{2+}$ activity was recorded in node 2. (*Bottom*) Representative $Ca^{2+}$ traces in node 2 in geometries with node/shaft width ratio

$\delta = \dfrac{d_0}{\mathrm{d_{shaft}}} = 1$ (red and gray), 2 (black) and 3 (blue), expressed as SNR (see Methods). Red and grey traces correspond to $Ca^{2+}$ signals in geometries with $\delta$=1 and a volume $V_1$=0.620 $\mu m^3$ and $V_1$=0.258 $\mu m^3$, respectively. Black and blue traces were recorded in geometries with $V_2$=0.263 $\mu m^3$ and $V_3$=0.195 $\mu m^3$, respectively. (*C*) Quantification of $Ca^{2+}$ signal characteristics depending on node/shaft width ratio $\delta$ and on process volume. Data are represented as mean ± STD, n=20 for each geometry. (*C1*) $Ca^{2+}$ peak probability does not vary with $\delta$ when $V_1$=0.258 $\mu m^3$ (p-value=0.22). (*C2*) Time to $1^{st}$ peak decreases with $\delta$ (p-value=0.0018 *). (*C3*) Peak amplitude does not vary with $\delta$ when $V_1$=0.258 $\mu m^3$ (p-value=0.16). (*C4*) Peak duration increases with $\delta$, for both $V_1$=0.620 $\mu m^3$ and $V_1$=0.258 $\mu m^3$ (***). (*C5*) The frequency of $IP_3R$ opening increases with $\delta$, for both $V_1$=0.620 $\mu m^3$ and $V_1$=0.258 $\mu m^3$ (***). (*D*) $Ca^{2+}$ residency time in nodes increases with $\delta$, for both $V_1$=0.620 $\mu m^3$ and $V_1$=0.258 $\mu m^3$ (***, n=300). The effect of node/shaft width ratio on each $Ca^{2+}$ signal characteristic was tested using one-way ANOVA. Significance is assigned by * for $p \leq 0.05$, ** for $p \leq 0.01$, *** for $p \leq 0.001$.

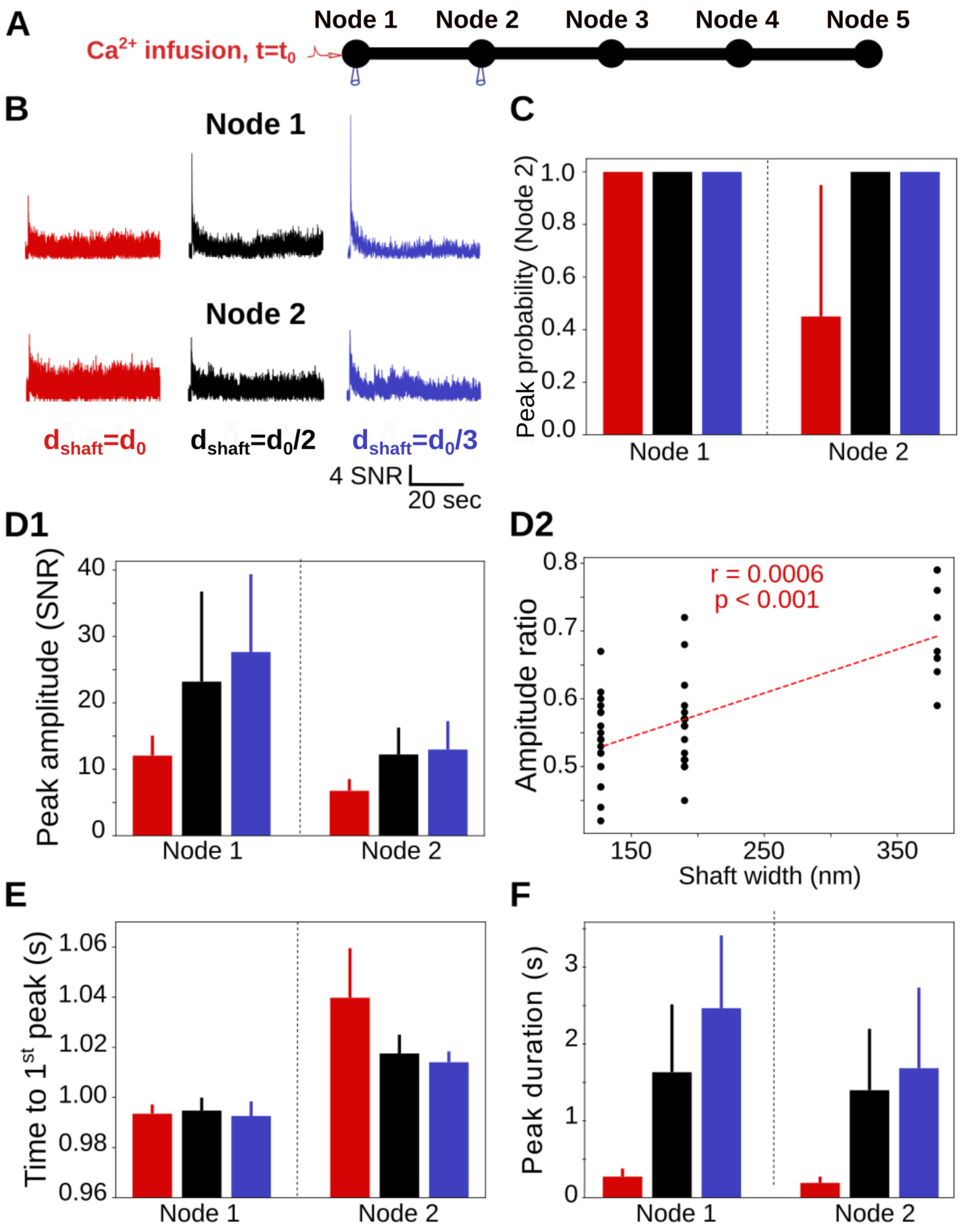


**Figure S6. Thin shafts enhance local spontaneous $Ca^{2+}$ activity.** (*A*) Neuronal stimulation protocol simulated for each geometry. 200 $Ca^{2+}$ ions were infused in Node 1 at time t=$t_0$, while $Ca^{2+}$ activity was recorded in nodes 1 and 2. (*B*) Representative $Ca^{2+}$ traces in node 1 (top) and node 2 (bottom) for $d_{shaft} = d_0$ (red), $\frac{d_0}{2}$ (black) and $\frac{d_0}{3}$ (blue), expressed as SNR (see Methods). (*C*) $Ca^{2+}$ peak probability increases when $d_{shaft}$ decreases in node 2 (***). Note that peak probability is 1 in node 1 as the injection of 200 $Ca^{2+}$ ions in node 1 results in a local Ca-GCaMP peak for all seed values tested. (*D1*) Peak amplitude increases when $d_{shaft}$ decreases in node 1 (***) and 2 (***). (*D2*) The ratio between peak amplitude in node 2 and in Node 1 increases with shaft width (Spearman r=0.0006, p-value < 0.001 ***). (*E*) Time to $1^{st}$ peak in node 2 increases with $d_{shaft}$ (***). Note that time to $1^{st}$ peak does not vary with $d_{shaft}$ in node 1 as it occurs quickly after $Ca^{2+}$ injection, through the binding of the injected $Ca^{2+}$ ions to GCaMP molecules. (*F*) Peak duration increases when $d_{shaft}$ decreases in node 1 (***) and 2 (***). Note that $Ca^{2+}$ peak characteristics were not statistically different between nodes 1 and 2. Data are represented as mean ± STD, n=20 for each geometry. The effect of $d_{shaft}$ on each $Ca^{2+}$ signal characteristic was tested using one-way ANOVA. Significance is assigned by * for $p \leq 0.05$, ** for $p \leq 0.01$, *** for $p \leq 0.001$.

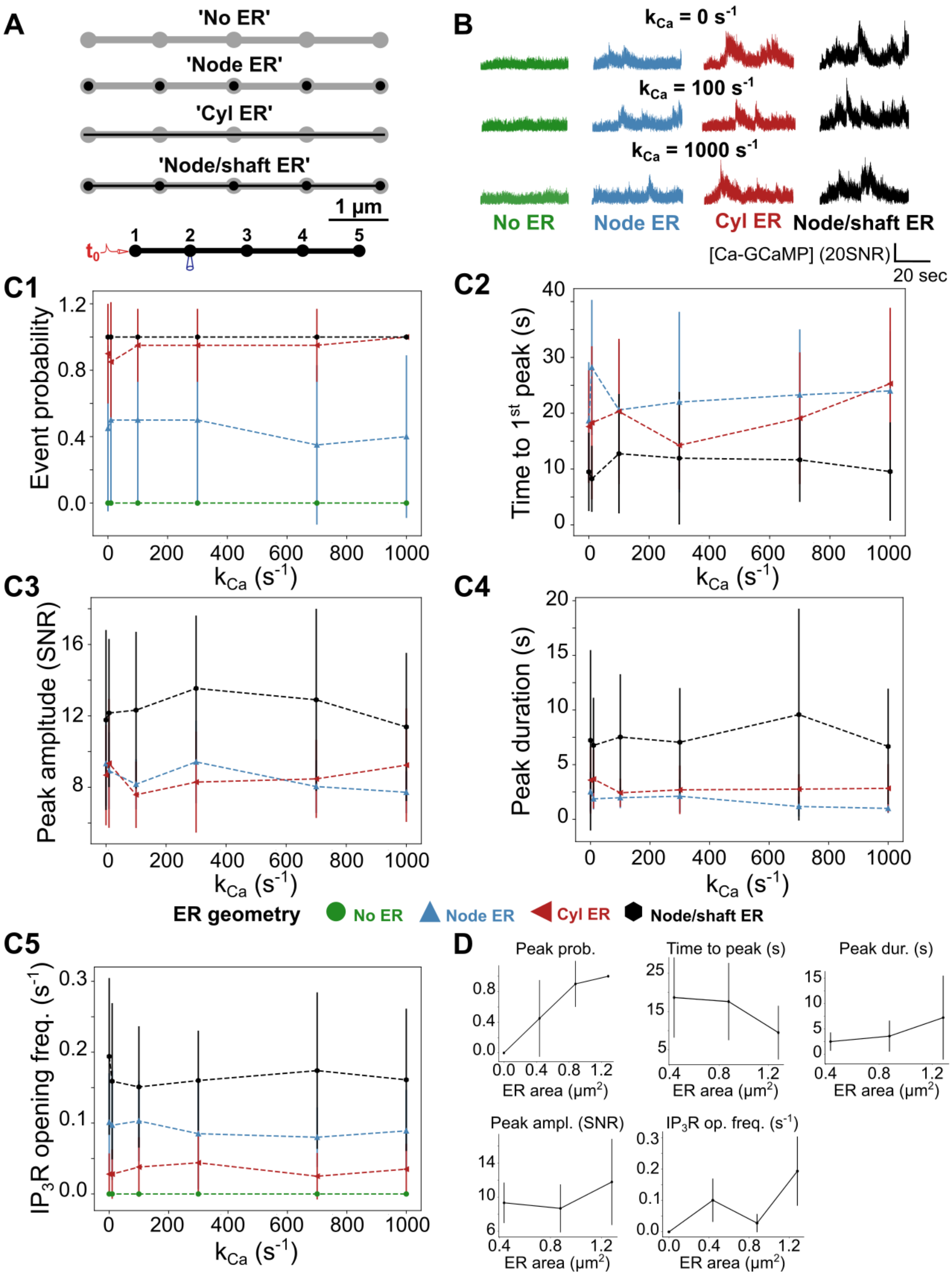


**Figure S7.** $Ca^{2+}$ **peak probability, amplitude and duration decrease with ER surface area.** (*A*) As the shape and distribution of the ER in fine processes have not been characterized in live tissue so far but are likely highly variable, simulations were performed in meshes with various

ER shapes: “No ER”, “Node ER”, “Cyl ER” and “Node/shaft ER”, in which there was no ER, discontinuous ER in nodes, cylindrical ER, or an ER consisting in node/shaft alternations, respectively. (*B*) (*Top*) Neuronal stimulation protocol simulated for each geometry: node 1 was stimulated at t=$t_0$=1s, while $Ca^{2+}$ activity was monitored in nodes 1 and 2. (*Bottom*) Representative $Ca^{2+}$ traces in node 2, in “No ER” (green), “Node ER” (blue), “Cyl ER” (red) and “Node/shaft ER” (black) geometries $d_{shaft} = \frac{d_0}{2}$, for $k_{Ca}$=0, 10, 100, 300, 700 and 1000 $s^{-1}$, expressed as SNR (see Methods). (*C*) Quantification of peak characteristics depending on ER geometry and $k_{Ca}$. $Ca^{2+}$ peak probability (***, *C1*), time to $1^{st}$ peak (***, *C2*), peak amplitude (***, *C3*), duration (***, *C4*) and the frequency of $IP_3R$ opening (***, *C5*) vary with ER morphology. Note that $Ca^{2+}$ peak characteristics did not vary with $k_{Ca}$. (*D*) Time to $1^{st}$ decreases as ER surface area increases (***) while $Ca^{2+}$ peak probability (***), amplitude (***) and duration (***) as well as the frequency of $IP_3R$ opening (***) increase as ER surface area increases, $k_{Ca}$=0 $s^{-1}$. Data are represented as mean ± STD, n=20 for each geometry. Significance is assigned by * for $p \leq 0.05$, ** for $p \leq 0.01$, *** for $p \leq 0.001$.

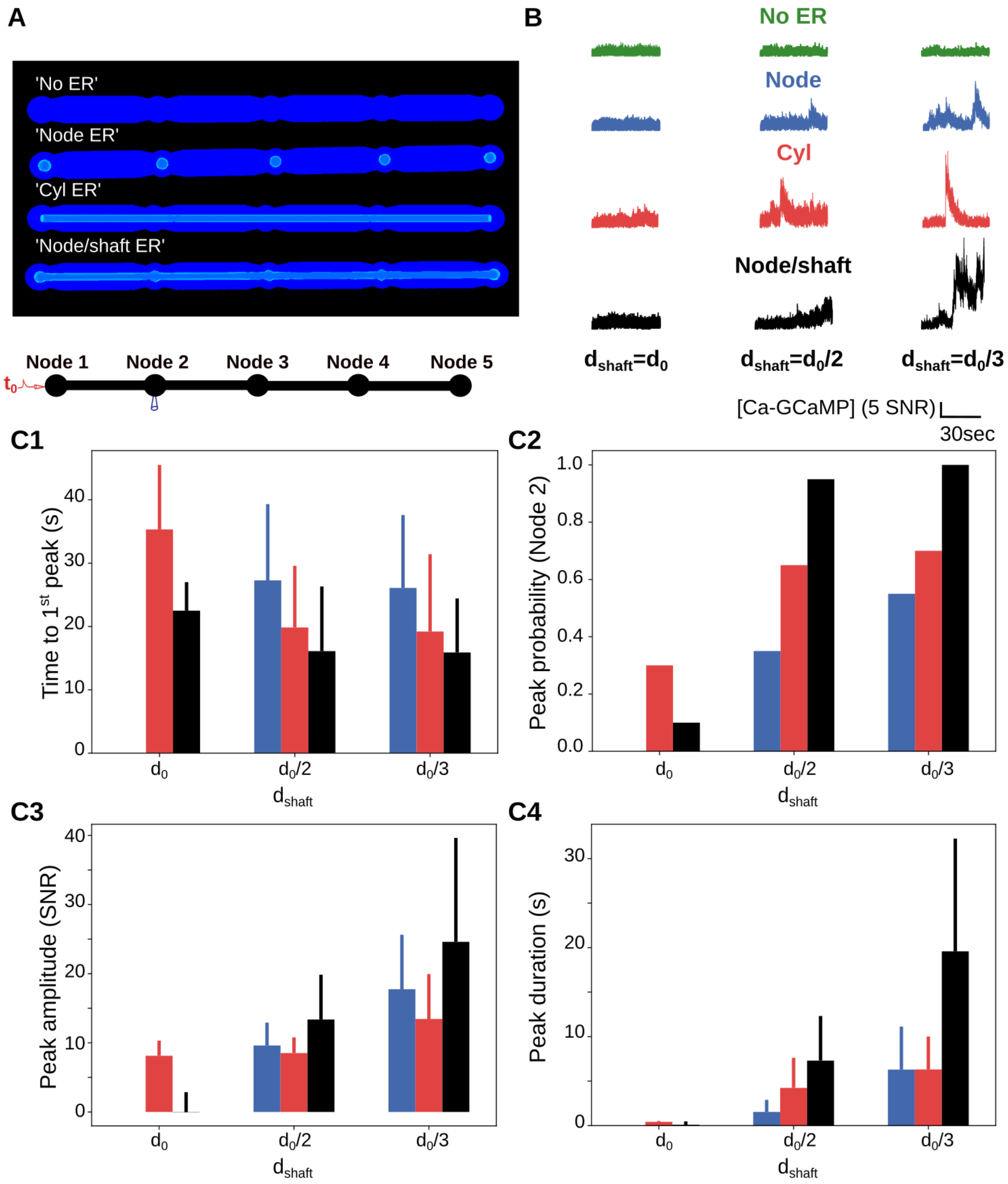


**Figure S8. Local $Ca^{2+}$ activity increases when shaft diameter decreases, for all ER geometries tested.** (*A*) Screenshots displaying the 3D meshes used to investigate the effect of ER geometry on $Ca^{2+}$ dynamics: "No ER", "Node ER", "Cyl ER" and "Node/shaft ER". (*B*) (*Top*) Neuronal stimulation protocol simulated for each geometry: node 1 was stimulated at t=$t_0$=1s ($k_{Ca}$=0 $s^{-1}$), while $Ca^{2+}$ activity was monitored in node 2. (*Bottom*) Representative $Ca^{2+}$ traces in node 2 for $d_{shaft}=d_0$ (left), $\frac{d_0}{2}$ (center) and $\frac{d_0}{3}$ (right), in "No ER" (green), "Node ER" (blue),

"Cyl ER" (red) and "Node/shaft ER" (black) geometries, expressed as SNR (see Methods). (*C*) Quantification of the effect of $d_{shaft}$ on $Ca^{2+}$ dynamics. Note that no bar is visible for simulations in "Node ER" $d_{shaft} = d_0$ geometries as no peak was detected. Results in "Node/shaft ER" geometries are presented in Fig3 but are added here (black bars) for comparison. (*C1*) Time to $1^{st}$ peak in node 2 does not vary with $d_{shaft}$ in "Node ER" simulations (p-value=0.85) but increases with $d_{shaft}$ in "Cyl ER" simulations (p-value= 0.021 *). (*C2*) $Ca^{2+}$ peak probability in node 2 increases when $d_{shaft}$ decreases in "Node ER" simulations (***) and in "Cyl ER" simulations (p-value=0.01 *). (*C3*) Peak amplitude increases when $d_{shaft}$ decreases in "Node ER" geometries (p-value=0.027 *) and in "Cyl ER" (p-value=0.01 *). (*C4*) Peak duration increases when $d_{shaft}$ decreases in "Node ER" simulations (p-value=0.028 *) and in "Cyl ER" simulations (p-value=0.001 **). Data are represented as mean ± STD, n=20, for each geometry. The effect of $d_{shaft}$ on each $Ca^{2+}$ signal characteristic was tested using one-way ANOVA. Significance is assigned by * for $p \leq 0.05$, ** for $p \leq 0.01$, *** for $p \leq 0.001$.

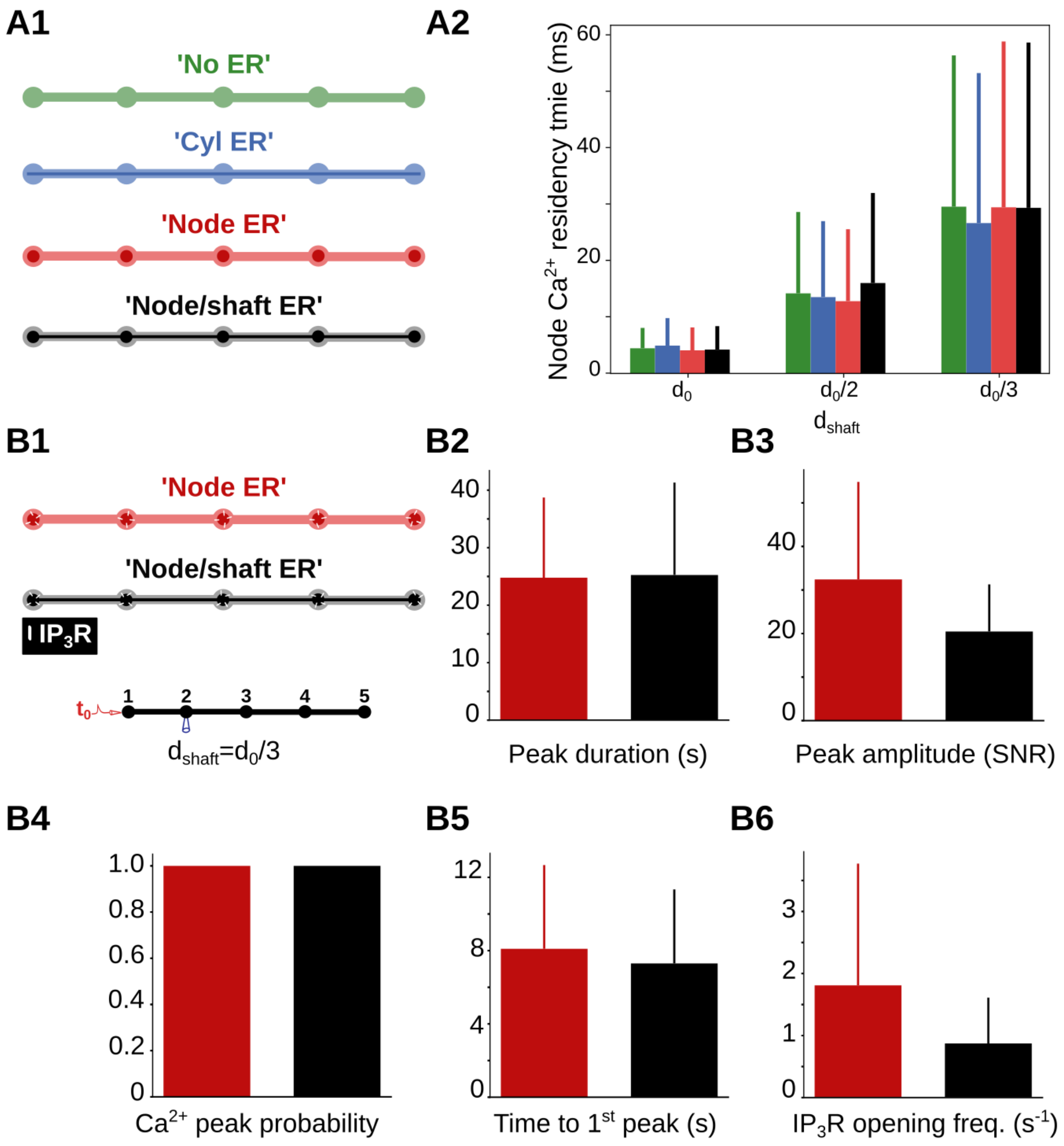


**Figure S9. Sensitivity study of the effect of ER morphology on $Ca^{2+}$ dynamics.** (A) Node $Ca^{2+}$ residency time is not influenced by ER morphology. (A1) One $Ca^{2+}$ ion was added in node 1 in geometries with different ER shapes: "No ER" (green), "Cyl ER" (blue), "Node ER" (red) and "Node/shaft ER" (black). (A2) For all ER shapes tested, $Ca^{2+}$ residency time in nodes increases when $d_{shaft}$ decreases. Node $Ca^{2+}$ residency time does not change depending on ER morphology, suggesting that cellular morphology has a greater influence on compartmentalization than ER morphology in the meshes used in this study. Data are represented as mean $\leq$ STD, n=300 for each geometry. (B) Sensitivity study of the effect of ER morphology on $Ca^{2+}$ dynamics with constant number of $IP_3Rs$. (B1) Simulations were performed in 'Node ER' (red) and "Node/shaft ER" (black) geometries with $IP_3R$ channels (white) located in nodes (Top), resulting in a constant number of IP3R channels for both ER shapes. Node 1 was stimulated at t=t0=1s ($k_{Ca}$=0 $s^{-1}$), while

$Ca^{2+}$ activity was monitored in node 2 (bottom). (B2-6) Quantification of $Ca^{2+}$ signals depending on ER morphology with constant number of $IP_3R$ in node 2, $d_{shaft} = \frac{d_0}{3}$. Data are represented as mean $\leq$ STD, n=20 for each geometry. $Ca^{2+}$ peak duration (B2), amplitude (B3), probability (B4) and delay (B5) do not vary with ER morphology when $IP_3R$ channels are located in nodes. (B6) $IP_3R$ opening frequency does not significantly differ between "Node ER" and "Node/shaft ER" geometries (p-value=0.06). Those results suggest that the decreased $Ca^{2+}$ activity in branchlets that contain discontinuous ER reported in Fig 4 mainly results from the decreased number of $IP_3R$ channels associated with the decreased ER surface area of "Node ER" compared to "Node/shaft ER" morphology.

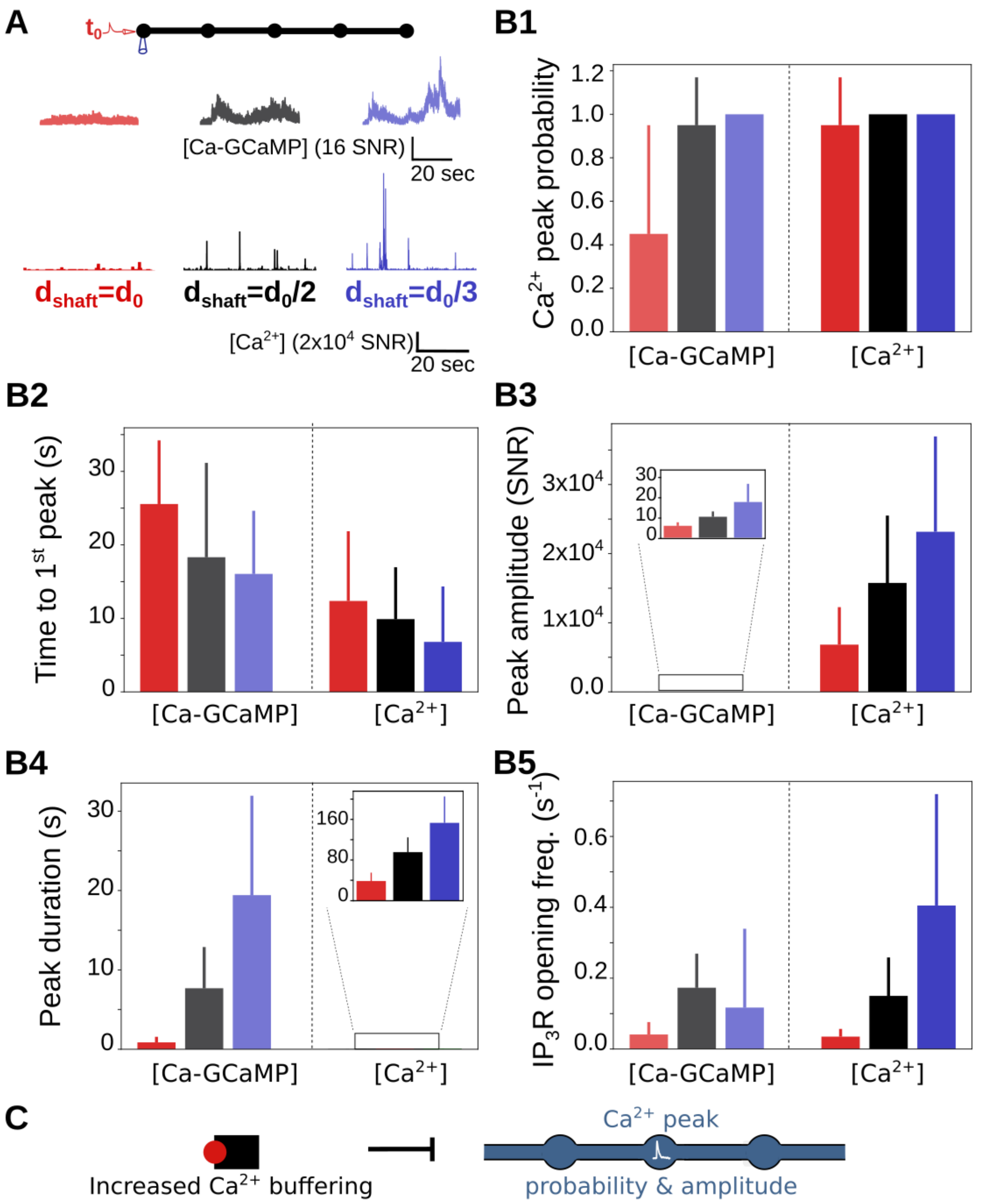


**Figure S10.** $Ca^{2+}$ **indicators alter** $Ca^{2+}$ **peak characteristics.** (*A*) In order to investigate free $Ca^{2+}$ peak probability, no GCaMP molecules were added to the model. (*Top*) At t=$t_0$=1s, node 1 was stimulated ($k_{Ca}$=0 $s^{-1}$), while $Ca^{2+}$ concentration was monitored in node 1. The geometry used for $d_{shaft}$ =$d_0$ corresponds to the geometry presented in Fig 1B. (*Middle*) Representative Ca-GCaMP traces in node 1, in simulations containing GCaMP molecules in geometries with shaft width $d_{shaft}$ = $d_0$ (light red), $\frac{d_0}{2}$ (grey) and $\frac{d_0}{3}$ (light blue). (*Bottom*) Representative free $Ca^{2+}$

traces in node 1, in the absence of GCaMP molecules, in geometries with shaft width $d_{\text{shaft}} = d_0$ (red), $\frac{d_0}{2}$ (black) and $\frac{d_0}{3}$ (blue). The amplitude of Ca-GCaMP and free $Ca^{2+}$ signals is expressed as SNR (see Methods). (*B*) Characteristics of free $Ca^{2+}$ signals depending on $d_{\text{shaft}}$. Ca-GCaMP signals measured in simulations from Fig 3 are presented for comparison. (*B1*) Free $Ca^{2+}$ peak probability does not vary with $d_{\text{shaft}}$ (p-value=0.22). Ca-GCaMP peak probability is lower than free $Ca^{2+}$ peak probability for $d_{\text{shaft}} = d_0$ (***). (*B2*) Time to $1^{st}$ free $Ca^{2+}$ peak increases with $d_{\text{shaft}}$ (***). Time to $1^{st}$ Ca-GCaMP peak is higher than time to $1^{st}$ free $Ca^{2+}$ peak, for any value of $d_{\text{shaft}}$ (***). (*B3*) Free $Ca^{2+}$ peak amplitude increases when $d_{\text{shaft}}$ decreases (***). Ca-GCaMP peak amplitude is lower than free $Ca^{2+}$ peak amplitude, for any value of $d_{\text{shaft}}$ (***). (*B4*) Free $Ca^{2+}$ peak duration increases when $d_{\text{shaft}}$ decreases (***). Ca-GCaMP peak duration is higher than free $Ca^{2+}$ peak duration, for any value of $d_{\text{shaft}}$ (***). (*B5*) The frequency of $IP_3R$ opening increases when $d_{\text{shaft}}$ decreases (***). (*C*) Schematic summarizing the main result from this figure: increased $Ca^{2+}$ buffering is associated with decreased $Ca^{2+}$ peak probability and amplitude. Data are represented as mean ± STD, n=20 for each geometry. The effect of $d_{\text{shaft}}$ on each free $Ca^{2+}$ signal characteristic was tested using one-way ANOVA. Significance is assigned by * for $p \leq 0.05$, ** for $p \leq 0.01$, *** for $p \leq 0.001$.

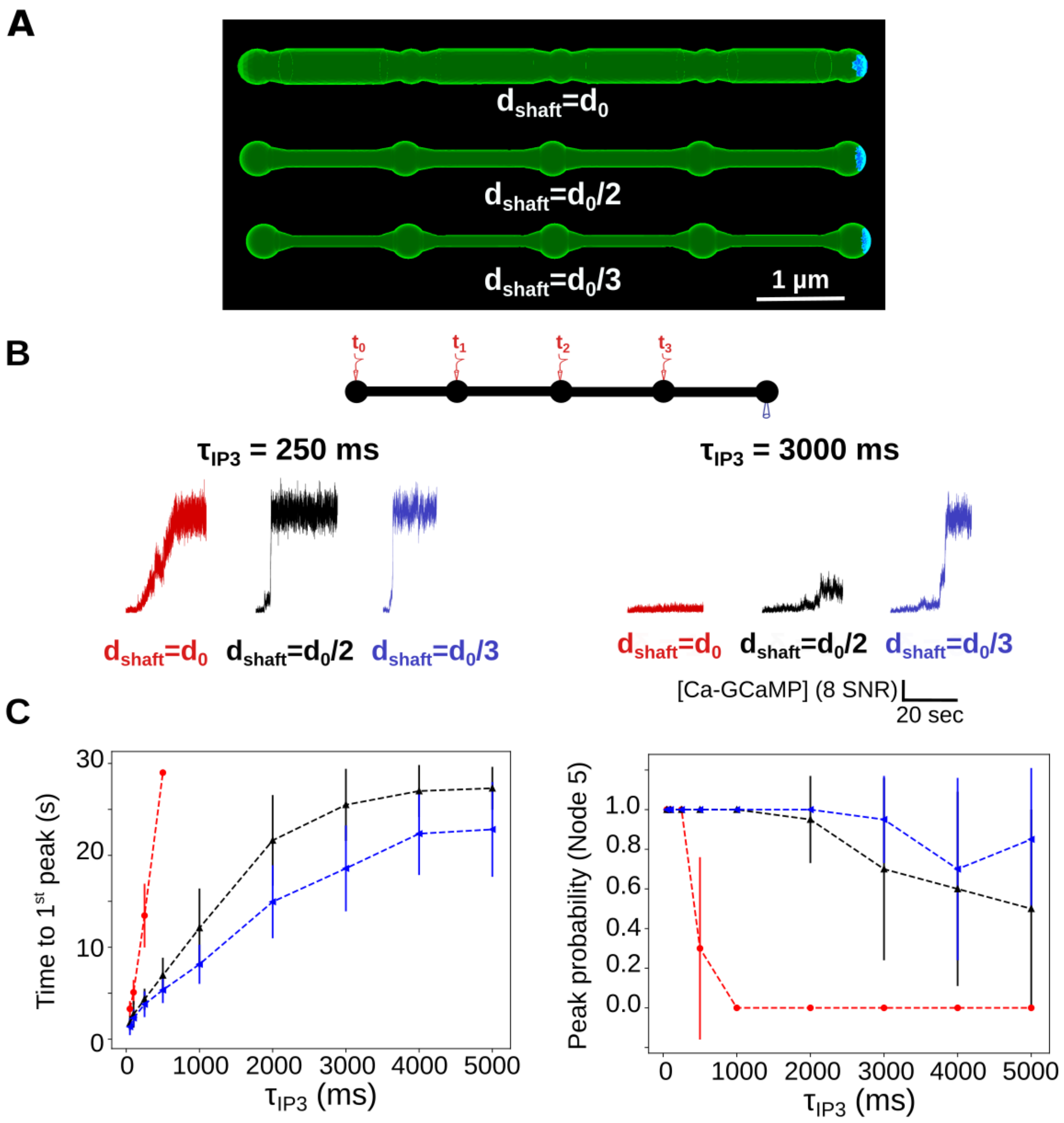


**Figure S11. Sensitivity study of the effect of boundary conditions on local** $Ca^{2+}$ **activity upon repeated neuronal stimuli.** (*A*) Simulations were performed in the "5nodes" geometries (Fig1). Boundary conditions were absorbing at the extremity of node 5 (blue) while the remaining boundaries were characterized by reflective boundary conditions (green). Those boundary conditions mimic the connection of the modeled astrocyte branchlet to the rest of the cell. (*B*) (*Top*) node 1 was stimulated at t=$t_0$=5s, node 2 at $t_0 + \tau_{\mathrm{IP3}}$, node 3 at $t_0 + 2\tau_{\mathrm{IP3}}$ and node 4 at $t_0 + 3\tau_{\mathrm{IP3}}$, $k_{Ca}$=0 $s^{-1}$, while $Ca^{2+}$ activity was monitored in node 5. (*Bottom*) Representative $Ca^{2+}$ traces for

$d_{shaft} = d_0$ (red), $\frac{d_0}{2}$ (black) and $\frac{d_0}{3}$ (blue), with $\tau_{IP3}$=250 ms (left) and 3000 ms (right), expressed as SNR (see Methods). (*C*) (*Left*) Time to $1^{st}$ peak in node 5 increases with $\tau_{IP3}$ for $d_{shaft} = d_0$ (***), $d_{shaft} = \frac{d_0}{2}$ (***) and $d_{shaft} = \frac{d_0}{3}$ (***). Time to $1^{st}$ peak is higher for $d_{shaft} = d_0$ compared to $d_{shaft} = \frac{d_0}{2}$ and $d_{shaft} = \frac{d_0}{3}$. (*Right*) $Ca^{2+}$ peak probability in node 5 is lower for $d_{shaft} = d_0$ compared to $d_{shaft} = \frac{d_0}{2}$ and $d_{shaft} = \frac{d_0}{3}$ and decreases as $\tau_{IP3}$ increases for $d_{shaft} = d_0$ (***) and $d_{shaft} = \frac{d_0}{2}$ (***) but not $d_{shaft} = \frac{d_0}{3}$ (p-value=0.054). Note that $Ca^{2+}$ peak probability was lower and time to $1^{st}$ peak higher compared to simulations with reflective boundary conditions. Data are represented as mean ± STD, n=20 for each value of $d_{shaft}$ and of $\tau_{IP3}$. Lines are guides for the eyes. The effect of $d_{shaft}$ on each $Ca^{2+}$ signal characteristic was tested using one-way ANOVA. Significance is assigned by * for $p \leq 0.05$, ** for $p \leq 0.01$, *** for $p \leq 0.001$.

### Movie S1

Movie presenting a simulation of the bleaching protocol, with $d_{shaft}=\frac{d_0}{2}$. Fluorescing ZsGreen molecules are in red and bleached, non-fluorescing, ZsGreen are in yellow. Molecule position was updated every 1 ms. The plot presents the variation of the number of fluorescing ZsGreen molecules with time in the central node. Note that, to facilitate visualization, here [ZsGreen] = 5 $\mu$M. Before the bleaching event, all ZsGreen proteins (red) fluoresce. At bleaching time, fluorescing molecules in the bleached region encounter conformational modifications that prevent them from fluorescing (yellow molecules). Consequently, the fluorescence level drops to $I_0$. Because of the diffusion of molecules in the cell, the fluorescence level in the central node then increases until it reaches a new baseline $I_{\text{inf}}$, after a recovery time $\tau$. Fluorescing ZsGreen molecules are in red and bleached, non-fluorescing, ZsGreen are in yellow.

### Movie S2

Movie presenting a simulation of the neuronal stimulation protocol, with $d_{shaft}=\frac{d_0}{2}$ and $Ca^{2+}$ influx rate at the plasma membrane $k_{Ca}$=0 $s^{-1}$. 100 $IP_3$ molecules are injected in Node 1 at t=$t_0$=0.1s, while the number of Ca-GCaMP, $IP_3$ and $IP_3R$ channels in the open state are monitored in nodes 1 and 2. Molecule position was updated every 0.1 ms. Ca-GCaMP molecules are in yellow, $IP_3$ in red and $IP_3R$ in blue. Note that the size of molecules in the movie was increased to facilitate visualization. Speed was increased 5-fold for visualization purposes.
